# Spiral NeuroString: High-Density Soft Bioelectronic Fibers for Multimodal Sensing and Stimulation

**DOI:** 10.1101/2023.10.02.560482

**Authors:** Muhammad Khatib, Eric Tianjiao Zhao, Shiyuan Wei, Alex Abramson, Estelle Spear Bishop, Chih-Hsin Chen, Anne-Laure Thomas, Chengyi Xu, Jaeho Park, Yeongjun Lee, Ryan Hamnett, Weilai Yu, Samuel E. Root, Lei Yuan, Dorine Chakhtoura, Kyun Kyu Kim, Donglai Zhong, Yuya Nishio, Chuanzhen Zhao, Can Wu, Yuanwen Jiang, Anqi Zhang, Jinxing Li, Weichen Wang, Fereshteh Salimi-Jazi, Talha A. Rafeeqi, Nofar Mintz Hemed, Jeffrey B.-H. Tok, Xiaoke Chen, Julia A. Kaltschmidt, James C.Y. Dunn, Zhenan Bao

## Abstract

Bioelectronic fibers hold promise for both research and clinical applications due to their compactness, ease of implantation, and ability to incorporate various functionalities such as sensing and stimulation. However, existing devices suffer from bulkiness, rigidity, limited functionality, and low density of active components. These limitations stem from the difficulty to incorporate many components on one-dimensional (1D) fiber devices due to the incompatibility of conventional microfabrication methods (e.g., photolithography) with curved, thin and long fiber structures. Herein, we introduce a fabrication approach, “spiral transformation″, to convert two-dimensional (2D) films containing microfabricated devices into 1D soft fibers. This approach allows for the creation of high density multimodal soft bioelectronic fibers, termed Spiral NeuroString (S-NeuroString), while enabling precise control over the longitudinal, angular, and radial positioning and distribution of the functional components. We show the utility of S-NeuroString for motility mapping, serotonin sensing, and tissue stimulation within the dynamic and soft gastrointestinal (GI) system, as well as for single-unit recordings in the brain. The described bioelectronic fibers hold great promises for next-generation multifunctional implantable electronics.

## Introduction

Biomedical devices with 1D geometries, such as surgical sutures, biopsy needles, endoscopes, guidewires, manometry probes, fiberscopes, and deep brain stimulation electrodes have been extensively used in the clinic for several decades (*1-6*). Compared to three-dimensional systems and 2D thin films, 1D devices are compact and can navigate through torturous paths. Therefore, 1D devices are favorable for deep penetration into tissues and body channels through minimally invasive implantation procedures. Furthermore, they can be readily retrieved and removed after completing their sensing or therapeutic functions (*1*).

Among the various examples of 1D devices, electronic fibers have gained considerable attention (*1, 7-9*). With recent advances in materials, fabrication, optics, and electronics, electronic fibers have acquired new functionalities, including sensing, actuation, tissue modulation, energy harvesting, and light emission (*10-17*). Despite their potential benefits, electronic fiber manufacturing remains challenging due to their incompatibility with conventional microfabrication techniques, which were originally developed for planar substrates. As a result, currently available fiber devices suffer from low density, limited functionality, and imprecise positioning of components. To date, approaches such as fiber spinning (*18-22*), 3D printing (*23, 24*), (micro)fluidics (*25, 26*), functionalization/coating of commercial fibers (*27, 28*), and thermal drawing (*29-32*), have been used to fabricate 1D fiber devices. Although electronic fibers prepared by thermal drawing can potentially host tens of channels and multiple functional components, the active region (which includes the sensing area) is typically located at the tip of the fiber, instead of distributed longitudinally. Additionally, it remains difficult to achieve high-density functions on fibers with precise control over device structure, position, and orientation. Examples of on-fiber lithography have been recently introduced to break the fiber’s longitudinal symmetry and to introduce micro-patterned components (*15, 33*). However, such procedures are still time-consuming and challenging for high-throughput production. In addition to the limitations mentioned above, the high temperature and high strain tolerance requirements for the thermal drawing process limit the choices of materials.

Herein, we introduce a versatile method to prepare high-density soft electronic fibers that can host a variety of functional components. We leverage state-of-the-art processes to microfabricate multifunctional sensors on a centimeter-scale 2D sheet and demonstrate that we can roll this into a monolithic micron-scale 1D fiber, called Spiral-NeuroString (S-NeuroString) (**Fig. 1A** and fig. S1). A variety of structural designs and advanced functionalities can be incorporated for various demanding biomedical applications, including sensing and stimulation in the highly dynamic and tortuous GI environment, and single neuron recording in the brain.

**Fig. 1.**
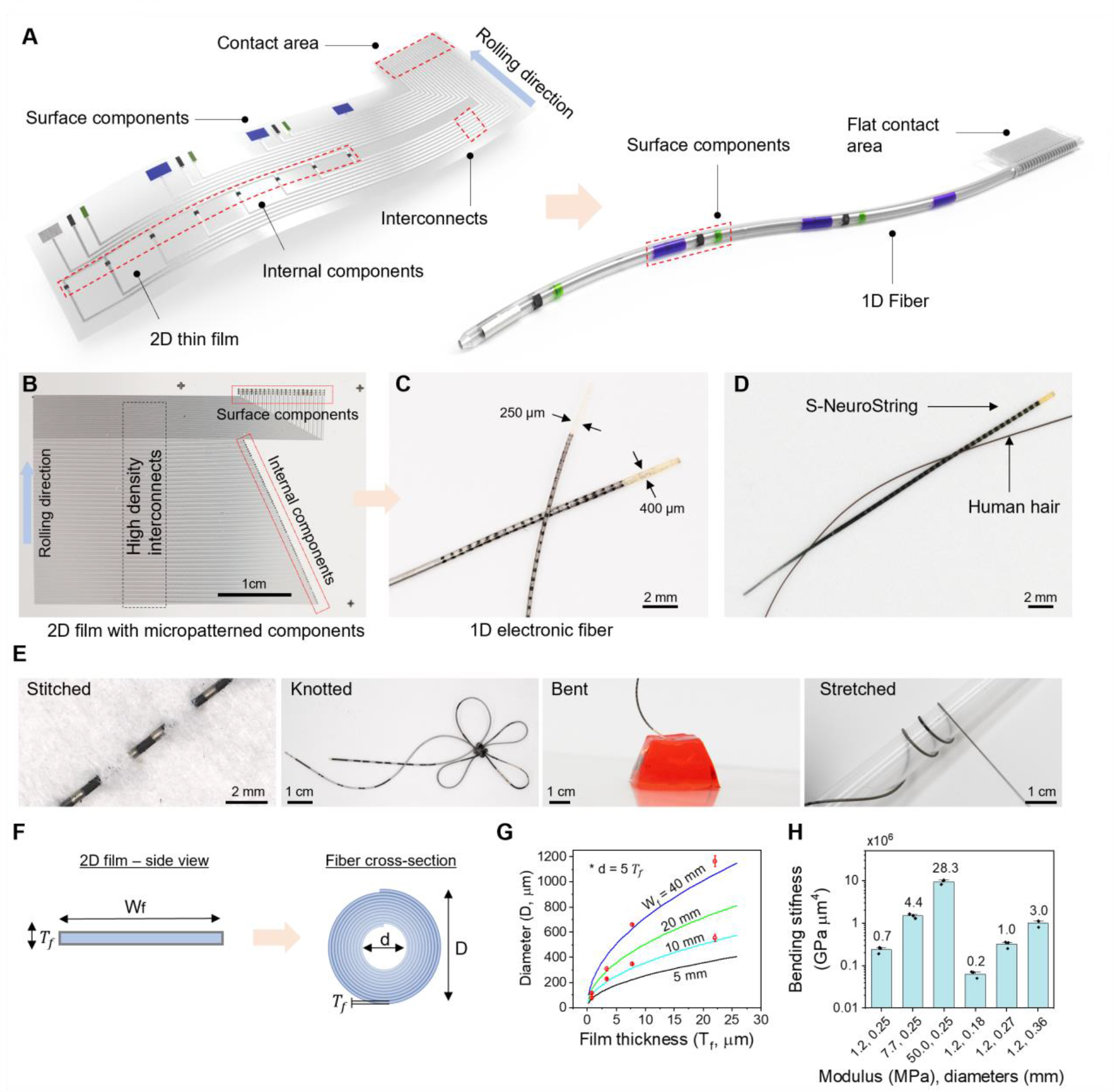
Fabrication of S-NeuroString. (**A**) Schematic illustration showing the transformation of a 2D film with micropatterned components into a 1D fiber. (**B**) A photograph of a 2D film with microfabricated devices used for preparing the thin high-density S-NeuroStrings shown in (C). (**C**) A photograph of two high-density S-NeuroStrings (250 and 400 µm in diameter) containing 150 sensors (81 pressure and 69 electrochemical sensors). The fibers were prepared from 3-cm-wide, 1.5- or 4.5-µm-thick substrates. A dramatic increase in density of devices per unit width follows the transformation from 2D to 1D, i.e., from a density of 150 components on 30 × 10 mm^2^ to 150 components per 0.25 × 10 mm^2^ projected area. (**D**) A photograph of S-NeuroString with 150 µm tip diameter next to a human hair. (**E**) Photographs showing the softness and flexibility of our bioelectronic fibers which can be stitched into a fabric, knotted, bent against soft gelatin, and stretched. (**F**) A schematic illustration of dimensions of 2D film and the corresponding 1D fiber. (**G**) Measured (red circles) and predicted fiber diameters as a function of 2D film thickness and length (as described in F). The measured diameters are slightly higher than the predicted diameters, due to slightly irregular rolling at the beginning of the process, which results in a larger internal diameter (n = 3 for each condition). (**H**) Bending stiffness calculated for fibers prepared from different types of SEBS (with different Young’s modulus: 1.2, 7.7, and 50.0 MPa) or varying 2D film thicknesses (which resulted in different fiber diameters: 0.18, 0.27, and 0.36 mm; n = 3).

## Results and Discussions

### Spiral transformation of 2D films into 1D S-NeuroString: design and fabrication

The rationale behind the development of bioelectronic fibers is that we can greatly increase the density of electronic components, compactness, and deformability (fig. S2). For the S-NeuroString preparation, we first performed microfabrication of desired electronic components on a 2D film using conventional deposition and patterning methods (**Fig. 1B**). Next, the 2D film was converted to the S-NeuroString using our spiral transformation approach (Movie S1, **Fig. 1C**). A S-NeuroString as thin as a human hair with 32 channels could be achieved. (**Fig. 1D)**. The S-NeuroString is soft, and can be stitched into fabric, knotted, bent, and stretched (**Fig. 1E**). The resulting fiber diameter depends on the width and thickness of the 2D film (**Fig. 1F,G**, Supplementary Section S1), while its stiffness can be controlled by the diameter or the modulus of the substrate material (**Fig. 1H &** fig. S3).

Notably, besides high-density devices that can be readily achieved by standard microfabrication on 2D films, the spiral transformation further boosts the density (per unit width) of components within the fiber by up to 2-3 orders of magnitude (fig. S4, Supplementary Section S2). For example, the 250 µm-diameter fiber shown in **Fig. 1C** was obtained from a 30 × 10 (sensing area only) × 0.0015 mm^3^ (W × L × T_f_) 2D film with 150 micropatterned sensing components (69 surface and 81 embedded) (**Fig. 1B**, fig. S5). The component density per unit width increased by 120 times, from 150/30 = 5 mm^-1^ to 150/0.25 = 600 mm^-1^.

The spiral transformation process can be divided into two main steps: (i) forming the core of the fiber (fig. S6A), and (ii) rolling the fiber (fig. S6B). The core is created by scraping against the edge of the 2D film, which collapses on itself to form a ‘fibrous’ structure which was then used to initiate the rolling process. Thin, soft, and highly self-adhesive films allowed for easier core formation (fig. S7). Alternatively, a predefined fiber core (e.g., metal and plastic wire, optic fiber, etc.) may be used to initiate the rolling process. This inner core can also be easily modified to provide additional functionalities (discussed next) and can also be used to tune the stiffness and the final diameter of the fiber. In the second step, the film was rolled until approaching the interconnect contact area to achieve the final fiber (fig. S8). Optimization of the rolling process was necessary to achieve densely packed fibers with intact shape and structure (fig. S6C). Thermoplastic styrene ethylene butylene styrene copolymer (SEBS) was employed as the main stretchable substrate due to its self-adhesive property (*34*), which is critical for both steps. While the adhesion between SEBS layers is sufficient to bond the rolled structure, we also incorporated an azide cross-linker (4 wt%), bis(6-((4-azido-2,3,5,6-tetrafluorobenzoyl)oxy)hexyl) decanedioate (*35*), to provide further stabilization by thermal or UV cross-linking. Other elastic materials such as styrene butadiene styrene copolymer (SBS) were also successfully used to prepare fibers.

The above versatile transformation approach allows us to create precise fibers with various device layouts (**Fig. 2A,B**, fig. S9 and Supplementary Section S3). For instance, some components can be positioned on the surface while others can be embedded inside the fiber based on their intended functions. In addition, the shape and structure of fibers can be controlled (fig. S10), and can readily incorporate additional functions, such as fluid delivery using hollow fibers or light transmission by integrating an optical fiber into the core (**Fig. 2C**, fig. S11). Additionally, this method can be extended to flexible substrates such as polyimide. We created electronic polyimide fibers containing multiple electrical resistors. The change in resistance after transformation was negligible, confirming the compatibility of the transformation process with non-stretchable substrates (**Fig. 2D**). However, unlike self-adhesive elastic materials, flexible substrates require an additional adhesive layer to help preserve its rolled-up structure.

**Fig. 2.**
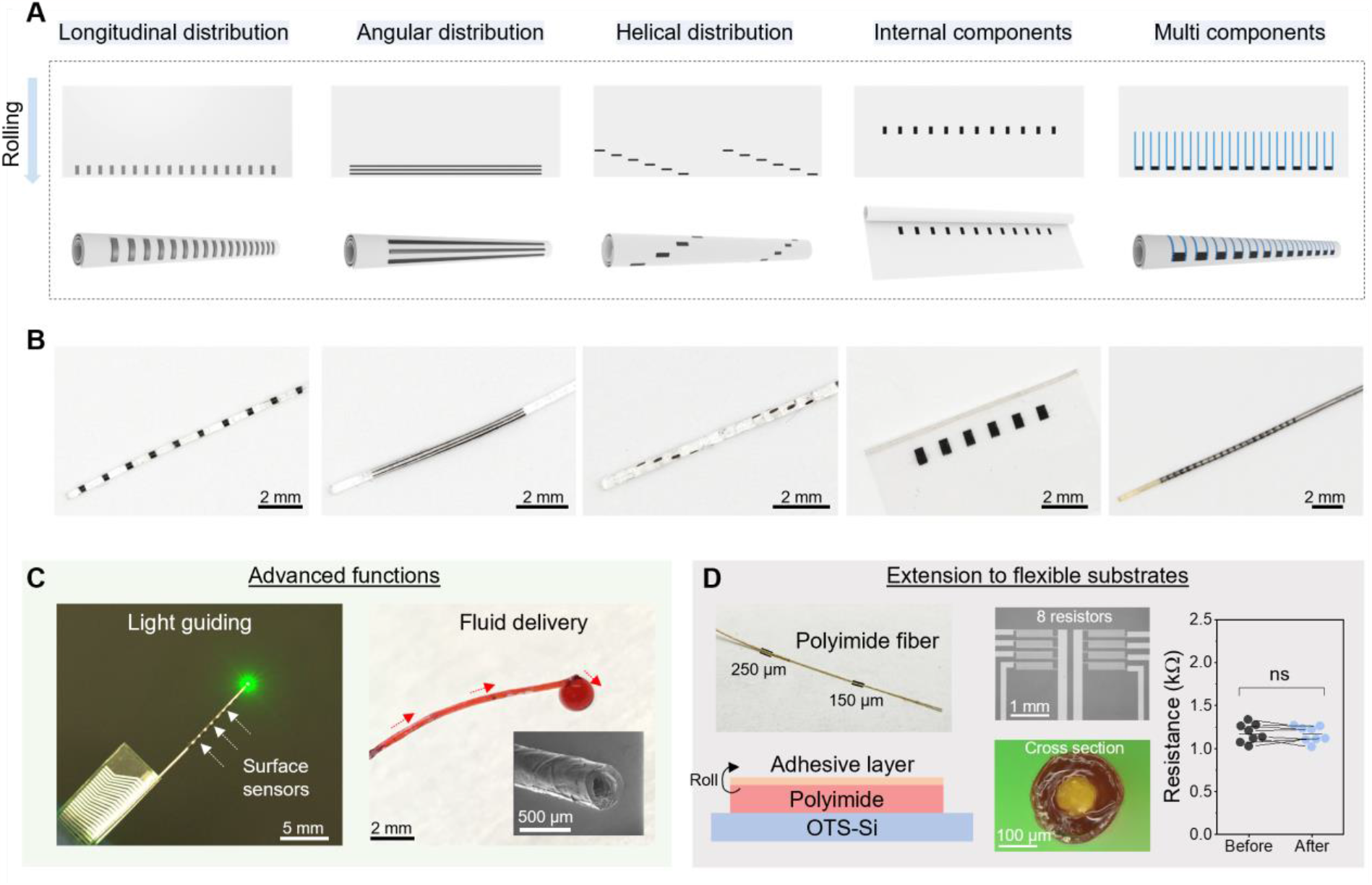
Versatility of structural design and advanced capabilities of the ″spiral transformation″. (**A**) Schematic illustration of 2D films with various layout designs and the corresponding fiber structure obtained after rolling. (**B**) Photos of the resulting fiber from the different designs shown in (A). (**C**) Advanced functions can be integrated, such as a light guiding optical fiber core (left) and a fluid delivery hollow fiber core (right). The hollow fiber (350 µm in diameter) became red due to the color of the liquid. (**D**) Extension to a flexible polyimide substrate (930 nm thick) with multiple micropatterned resistors. The measured resistance before and after the rolling process showed little change (n = 8 resistors). Paired, two-tailed Student’s t-test.

### S-NeuroString for deployment in the gut

The GI tract is challenging to access due to its tortuous structure and constant peristalsis (*36-38*). While smart-pill sensors have been developed, they cannot simultaneously perform sensing or actuation over long distances (*39-41*). In this work, since the diameter of our 1D soft S-NeuroString is order of magnitudes lower than smart pills, we leverage its geometrical and mechanical compliance to enable sensing and modulation in the GI tract over multi-centimeter length scales. Our multifunctional GI S-NeuroString includes functional components such as strain/pressure and serotonin sensors as well as electrical stimulation electrodes (fig. S12). Eutectic gallium indium (EGaIn) was used as the conductive interconnects due to its high conductivity and stretchability (fig. S13). The sensing components were all prepared from laser-induced graphene (LIG, fig. S14) (*37, 42*). A double network of poly(3,4-ethylenedioxythiophene) polystyrene sulfonate (PEDOT-PSS) and poly(ethylene glycol)-block-poly(propylene glycol)-block-poly(ethylene glycol) (PEG-PPG-PEG) was used for preparing the electrical stimulation electrodes due to its overall high conductivity, low interfacial impedance, and good stretchability (*35, 43*).

All electrical components were first tested and characterized *in-vitro*. The pressure/strain sensors were based on simple LIG-based piezoresistors that change resistance as a function of applied tensile and/or compressive forces. We observed sensitive and reversible responses for forces between 0.1 – 2 N (fig. S15). S-NeuroString provided excellent longitudinal mapping of forces, detection of progressive forces, as well as information on curvatures at various locations along the fiber (fig. S15-18), all of which are relevant to studying the biomechanics and motility patterns of the GI tissue. For serotonin sensing, LIG-based electrodes were used with chronoamperometry or cyclic voltammetry according to our previous report (*37*). Good sensitivity (0.332 nA nM^-1^) and an extrapolated detection limit of ∼24 nM were obtained (fig. S19). For electrical stimulation, the low impedance of the PEDOT-PSS-based electrodes allowed us to achieve high stimulation currents (< 5 mA) under low voltages (< 4V, fig. S20). Finally, we could add pH sensing to our S-NeuroString (fig. S21).

Our S-NeuroString is a suitable candidate for interfacing with the GI tract due to the fiber’s high softness and isotropic deformability. We observed negligible motility disruption after inserting a fiber (300 µm in diameter and ∼ 5 MPa modulus) into an isolated mouse colon (*ex-vivo*, fig. S22). Moreover, the presence of the fiber inside the colon of the awake mice (*in-vivo*) did not show any adverse effects on their behavior. In contrast, insertion of polyimide (previously used for motility sensing devices (*44*), having a modulus of > 1GPa) rectangular films led to significantly diminished movement and clear hematochezia (i.e., blood in stool pellets, fig. S23A-D). These findings were further supported by our obtained histological images that revealed significant disruptions to the native tissue architecture caused by the flexible substrate but not by our soft fibers. fig. S23E).

To highlight the multifunctionality and the spatial mapping capabilities of the S-NeuroString, we next demonstrated simultaneous *ex-vivo* colonic motility sensing and stimulation in mice. The S-NeuroString with distributed pressure sensors was able to detect temporal and spatial motility signals with high precision (fig. S24A,B). Notably, natural spatially progressing contraction events were observed and detected (fig. S24C-G and Movie S2). Using surface electrodes on S-NeuroString to electrically stimulate the GI tissue, according to a previously reported biphasic stimulation method (*45*), we observed an increase in motility events from the responses of distributed piezoresistive sensors (fig. S25A). This resulted in a significant increase in motility index (defined as the average area under recorded motility peaks per minute), as calculated from the sensory output of the S-NeuroString (fig. S25B-C). Video analysis also confirmed larger changes in colonic diameter as well as higher frequency of events detected under stimulation, including both slow waves and colonic migrating motor complexes (CMMCs) (fig. S25D-E). Additional colonic motility tests were performed using the S-NeuroString *in-vivo* in anesthetized mice (fig. S26).

To validate the potential application of our S-NeuroString in larger animal models, we tested both sensing and modulation functions in the small intestine of swine. First, for natural motility sensing, S-NeuroString with a 10 cm sensing length was inserted in the small intestine via laparotomy (**Fig. 3A,B**). Despite the complexity and fluctuation of the *in-vivo* GI environment, we managed to observe and record natural motility patterns multiple times (**Fig. 3C**, fig. S27 and Movie S3). Notably, the S-NeuroString (300 µm in diameter) was ∼50-fold smaller than the diameter of the intestine but it was still able to provide reliable sensing of motility signals. Next, we designed a bifunctional S-NeuroString for closed loop motility sensing and stimulation (**Fig. 3D,E**). The response of pressure sensors to contractions induced by stimulation is shown in **Fig. 3F,G**. A threshold of 2 V was required to induce noticeable contraction signals. This bifunctionality allowed us to perform a programmable stimulation (**Fig. 3H**). Distributed stimulation electrodes were used to stimulate specific parts of the tissue and to program the progression of contractions (Movie S4) -a process that resembles the naturally-generated peristalsis. At the same time, distributed pressure sensors provided feedback about the location and progress of contractions (**Fig 3I**). This system can be potentially used for real-time optimization and personalization of electrical stimulation conditions.

**Fig. 3.**
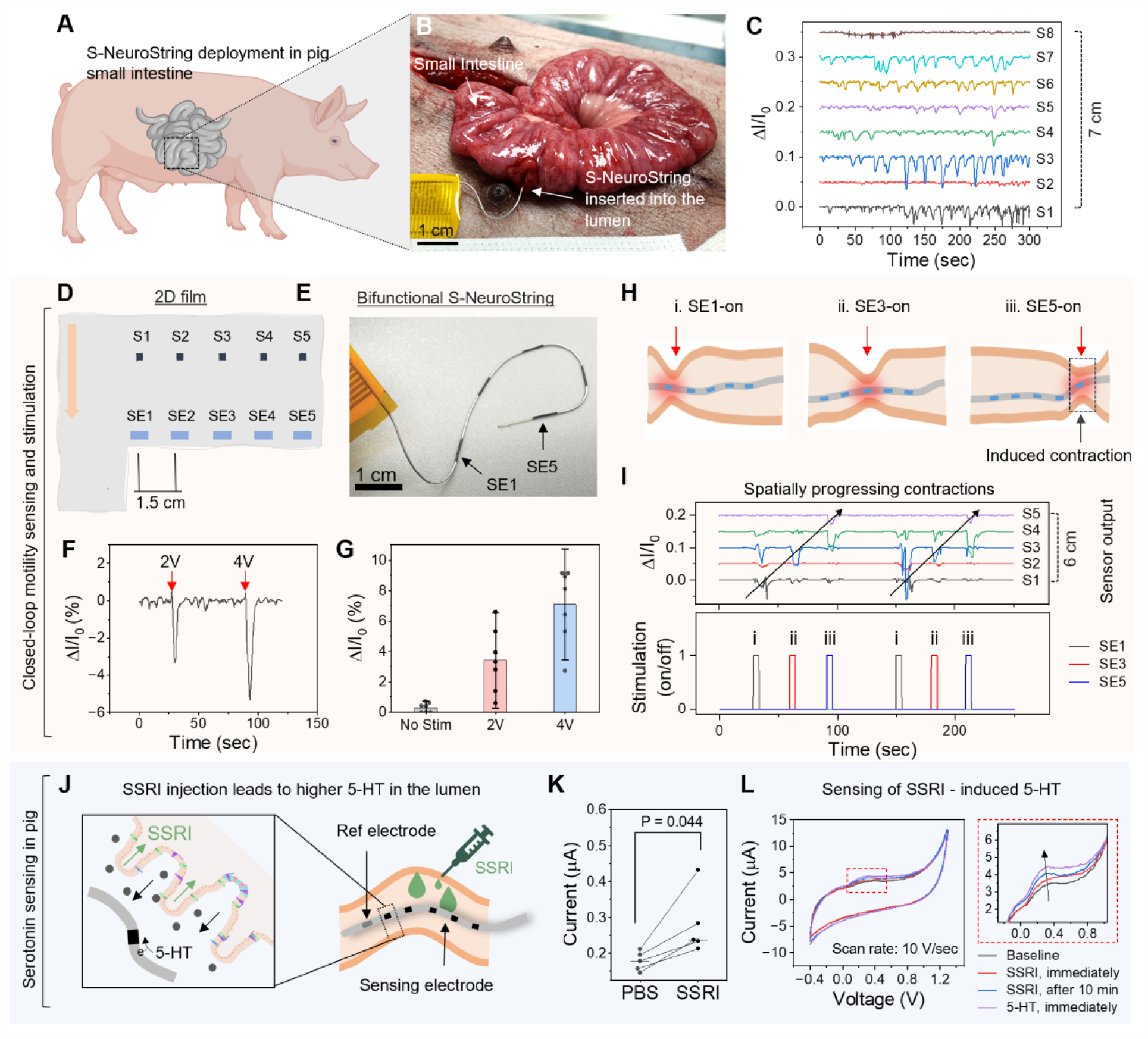
S-NeuroString for sensing and stimulation in the GI tract of pigs. (**A**)**-**(**B**), Schematic and photographic illustrations showing the deployment of an S-NeuroString into the small intestine of pig, which was used for monitoring and modulating GI signals. (**C**) Recording of natural motility in the small intestine using a S-NeuroString with 8 pressure sensors (S1-S8). All traces are normalized and shifted for clarity (see methods). (**D**) Schematic illustration of the 2D design of a bifunctional S-NeuroString used for simultaneous motility sensing (S1-S5) and stimulation (SE1-SE5). (**E**) A photograph showing the bifunctional S-NeuroString obtained from the 2D design shown in (D). (**F**) Pressure sensor output in response to stimulation at 2 and 4 V. (**G**) Summary of the response obtained under stimulation at different voltages (n = 7 stimulation experiments, 2 S-NeuroStrings, 2 pigs). The bars represent the mean, and the error bars represent the standard deviation. The results in (F) and (G) were obtained from sensors and stimulation electrodes that are located at the same point. (**H**) Schematic illustration showing the use of distributed stimulation electrodes to induce programmable contraction events over long intestinal segments. (**I**) Demonstration of the programmable stimulation. Stimulation of the pig intestine at different locations generates progressive contractions similar to natural motility patterns. Distributed pressure sensors provide feedback about the location and progress of contractions. (**J**) Schematic illustration showing the use of SSRI injection (0.5 ml of 10 µM) to induce serotonin release inside the intestine. (**K**) Drug-induced luminal serotonin concentration change using an SSRI solution and PBS as control (n = 5 experiments, 3 S-NeuroStrings, 3 pigs). The short horizontal lines on the graph represent the means. Each point is the average of 5 electrochemical sensors as shown in fig. S28. Paired, two-tailed Student’s t-test. (**L**) Electrochemical sensing of SSRI-induced serotonin using cycling voltammetry.

We next demonstrated the use of S-NeuroString for electrochemical sensing of serotonin in the small intestine of swine (**Fig 3J)**. Using amperometry, changes in serotonin concentrations could be detected following the injection of selective serotonin reuptake inhibitor (SSRI) (**Fig 3K**, fig. S28). Cyclic voltammetry was used to verify the existence of serotonin peaks (**Fig 3L**).

### High Density S-NeuroString for Single Neuron Recording

Although elastomeric substrates are used extensively for neural interfaces due to their stretchability and conformability matching that of native tissue, previous studies have been primarily limited to recording population level activities (*35, 46-49*). Single-unit neuronal recordings impose more stringent requirements on electrical characteristics, which includes: (i) the recording site needs low interfacial impedance, small size, and high-density to detect and isolate single unit activity, (ii) adjacent electrodes should have a low cross-coupled impedance to minimize crosstalk, (iii) the dielectric has to provide a high shunt impedance to minimize attenuation (*50-53*). Hence, we next examined whether we could reconfigure our high-density S-NeuroString to meet these above demanding requirements.

First, we developed a bi-layer encapsulation strategy with photopatternable crosslinked SBS (3.2 µm thick) to reduce the likelihood of pinholes exposing the conductive traces. This resulted in a stable and high impedance of ∼ 1.7 MΩ at 1 kHz (for a 0.5 × 5 mm^2^ area of encapsulated electrode) across a period of 1 month in PBS (fig. S29). For the microelectrodes, we used a double network of PEDOT-PSS and PEG-PPG-PEG. We developed an etching process enabling the patterning of micron-scale electrodes and confirmed their low interfacial impedances (10.4 kΩ at 1 kHz, for a 25 × 25 µm^2^ electrode area) and stability across a period of 1 month (fig. S30).

Next, we evaluated S-NeuroString’s ability to record neuronal activity in awake and moving mice. We designed and fabricated a 32-channel 2D film that was then transformed into a 150-μm-diameter neural S-NeuroString (**Fig. 4 A,B** and fig. S31). The S-NeuroString microelectrodes were helically distributed across a 1.6 mm length in groups of tetrodes with 40 μm pitch (fig. S32). After transformation, the interfacial impedance of S-NeuroString microelectrodes showed good uniformity (**Fig. 4C**). Compared to existing microelectrode arrays used for single neuron recording, S-NeuroString’s Young’s modulus was a few orders of magnitudes lower (**Fig. 4D**). We next implanted S-NeuroStrings into multiple regions of the brain with an ultrasonic micron-scale vibration inserter to minimize the acute insertion trauma, and recorded neural signals in awake mice (**Fig. 4E)**. We observed spontaneous spiking activities (**Fig. 4F**), as early as two days post implantation (fig. S33). Using Kilosort3, a commonly used spike sorting pipeline, activities of individual neurons could then be identified (**Fig. 4G**). Due to the high electrode density, single unit waveforms were detected across multiple electrodes (**Fig. 4G**) enabling good isolation, as indicated by the low fraction of interspike interval (ISI) violations (<0.3) (**Fig. 4H** and fig. S34), high signal-to-noise ratio (SNR), and a high degree of separation in Uniform Manifold Approximation and Projection (UMAP) space (**Fig. 4I** and fig. S34). Finally, using immunohistochemistry, we showed that an S-NeuroString with 150-µm diameter had a minimal body response comparable to other reported fibers of similar diameters (fig. S35) (*36, 54*).

**Fig. 4.**
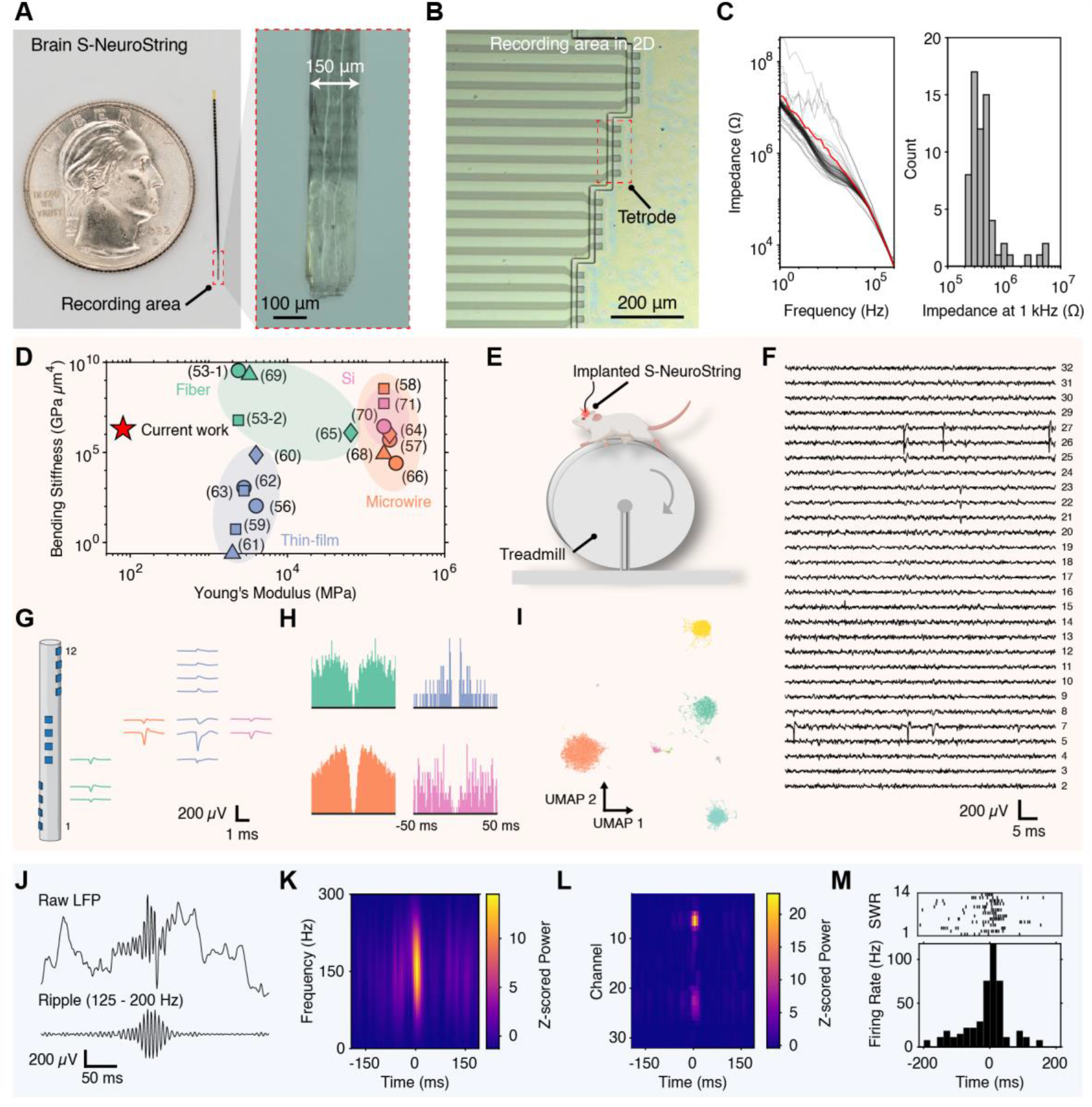
Recording of single unit activity and sharp wave ripples in awake and moving mice. (**A**) A photograph of the neural S-NeuroString used for recording in the brain. The inset shows a microscopy image of the recording tip. (**B**) A microscopy of the recording area of neural S-NeuroString, showing tetrodes of PEDOT-based microelectrode. **(C)** Impedance measurements of the PEDOT-based microelectrodes in saline solution (n = 64 electrodes from 2 fibers), showing a yield of > 90% for impedance < 1MΩ at 1kHz after fabrication and transformation. (**D**) Comparison of mechanical properties between S-NeuroString and existing microelectrode array-based devices used for single neuron recording. (**E**) A schematic of the behavioral rig used for recording neural activity in awake and moving mice. (**F**) A waterfall plot of the band-pass filtered data (300 – 7000 Hz) across 32 electrodes showing spontaneous spiking activity a week after S-NeuroString implantation into the hippocampus. (**G**) Schematic of the fiber with the corresponding spike-triggered averages, i.e., the footprint, of individual neurons adjacent to their respective electrodes. The high-density tetrodes enable multiple electrodes to detect the waveform of individual neurons. (**H**) Autocorrelograms of the respective neurons as shown in **f** with clear refractory periods indicative of isolated single neurons. (**I**) Waveforms plotted in UMAP space indicate a high degree of separation between individual clusters. (**J**) Raw and bandpass filtered traces of a representative SWR detected in the hippocampus. (**K**) Spectrogram showing an increase of power centered around 160 Hz, (**L**) Spatial representation of the SWR spectral power at 160 Hz across multiple electrodes revealing stronger engagement in CA1. **(M)** A representative neuron exhibiting elevated firing rates associated with the SWR events, as shown by the raster plot and peristimulus time histogram (PSTH).

With the S-NeuroString implanted into the hippocampus in mice, we confirmed that the probes were able to detect population level activity in the form of sharp wave ripples (SWR), short oscillatory bursts between 125 – 200 Hz (**Fig. 4J**) which are associated with memory consolidation and memory retrieval (*55*). We observed characteristic electrophysiological signatures such as the co-occurrence of CA1 and CA3 ripples, as well as an inverse of LFP polarity with electrode depth (fig. S36). Moreover, the spectrogram of individual channels exhibited a stereotypical increase in power (**Fig. 4K**) (*56*), with stronger engagement observed particularly in the CA1 region (**Fig. 4L**). Furthermore, several neurons exhibited elevated firing rates associated with SWR events (**Fig. 4M**, fig. S36).

## Discussion

We describe a strategy to fabricate high-density multimodal soft bioelectronic fibers capable of sensing and stimulation. Our approach takes advantage of the ease of microfabrication and the large area available on 2D films to incorporate circuit components and subsequently roll them into 1D fibers using a spiral transformation. Our developed S-NeuroString offers a substantial improvement in the density of functional components per unit width of the final device structure as compared to planar flat devices. The versatility of the approach is reflected in thin electronic fibers with control over the devices’ angular, radial, and longitudinal positions. Furthermore, as compared to existing electronic fibers, S-NeuroString not only exhibits much lower softness but also increased sensing units, density, and functionalities (summarized in Table S1). Moreover, our GI S-NeuroString exhibits numerous advantages over other sensing probes currently available for the GI tract, such as high resolution and density, compatibility with conventional microfabrication, and multimodality, as outlined in Table S2. In the context of neural recording, S-NeuroString shows multiple advantages when compared to existing microelectrode arrays with multiple recording channels (Table S3). Notably, for brain studies, S-NeuroStrings that combine light delivery for optogenetics, electrochemical sensors for neurotransmitters, and single neuron recording could be developed in the future using our transformation process. While the focus of our various demonstrations has primarily been on in-body implantation, our findings herein hold significant potential for a wide range of applications, including smart fabrics, textiles, wearables, and soft robotics.

## Acknowledgments

This work was partly supported by the Wu Tsai Neurosciences (Big Ideas in Neuroscience), Maternal and Child Health Research Institute at Stanford, CZ Biohub-San Francisco. Part of this work was performed at the Stanford Nano Shared Facilities (SNSF) and Stanford Nanofabrication (SNF). Part of this work was performed at Stanford Cell Sciences Imaging Facility (CSIF), which is funded by NIH ORIP funding: 1 S10 OD032300-01. We thank M. Sosa and E.A. Jones for insightful discussions on hippocampus electrophysiology. MK acknowledges postdoctoral fellowship support from the Fulbright Foundation and Israeli Council for Higher Education (VATAT). ETZ acknowledges support from the Bio-X Stanford Interdisciplinary Fellowship and Croucher Scholarship. CZ acknowledges the funding from an F32 fellowship from the National Institute of Biomedical Imaging and Bioengineering of the National Institutes of Health (F32EB034156). A.A. acknowledges funding from an NIH F32 fellowship (grant 1F32EB029787). ZB is a CZ BioHub investigator. XC and ZB acknowledge support from the Tianqiao and Chrissy Chen Institute (TCCI).

During the course of this work, we discovered similar approach on using rolling to prepare soft bioelectronic fiber is being pursued by Zhiyuan Liu Group in Shenzhen Institutes of Advanced Technology (SIAT), Chinese Academy of Sciences (CAS). We jointly decided to post our work simultaneously in BioRxiv. The work from Liu Group is entitled “Rolling 2D Bioelectronic film into 1D: a Suturable Long-term Implantable Soft Microfiber” by R. Xie, Q. Yu, D. Li and Z. Liu et al.

## Author contributions

M.K. and Z.B. conceived the project and designed the experiments. M.K. designed, and fabricated the devices, performed the mechanical, electrical and electrochemical measurements. D.C. helped in the characterization of chemical and pressure sensors. W.Y. performed the SEM. C.X. performed Raman characterization, surface roughness analysis of LIG, and design of photographic illustrations. J.P. helped in the preparation of connections and wiring parts used for the measurements, and design of schematic illustrations. Y.L., D.Z., K.K.K., Y.N., Y.J., C.W. and W. W. helped with the development of the materials and fabrication processes. S. E. R. captured digital microscopy images of fibers. J. L. advised the preparation and testing of the serotonin electrochemical sensors. M.K., R.H., and E.S. carried out all the *ex vivo* experiments. E.S. analyzed the *ex-vivo* data. A.A. and M.K. designed, performed, and analyzed the colonic biocompatibility tests and *in-vivo* colonic recording in anesthetized mice. M.K., C.-H. C., A.-L.T., F.S.-J. and T. A. R. performed the anesthetized pig experiments for the gut-related applications. M.K., E.T.Z. and S.W. designed the neural recording experiments with help from L.Y. and N.H.. M.K. E.T.Z. and S.W. performed all the neural recording experiments. E.T.Z. analyzed the neural recording data. S.W. and M.K. performed the brain histology experiments. S.W. analyzed the brain histology data. A. Z., helped with the histology study. A.A., E.T.Z., S.W., and C.Z. prepared the different mice protocols used in this study. A.-L.T. and J.C.Y.D. prepared the pig protocol. X.C. contributed to the neural recording study. J.A.K. contributed to the *ex-vivo* gastrointestinal studies. M.K. E.T.Z., S.W., J.C.Y.D., and Z.B. prepared the manuscript. All authors commented on the manuscript and gave approval to the final version. Z.B. and J.D. directed the project.

## Competing interests

Stanford University has filed a patent application related to this technology. The patent application number is 63/528846. A.A. is a co-inventor on a patent application describing ingestible devices for electrical stimulation. A full list of A.A.’s competing interests can be found here https://www.abramsonlab.com/aa-conflicts-of-interest.

## Supplementary Materials

Materials and Methods Figs. S1 to S35

Tables S1 to S3

Supplementary Section S1 to S3

References(*1-54*)

Movies S1 to S4

