## Supplementary Materials for "Spiral NeuroString: High-Density Soft Bioelectronic Fibers for Multimodal Sensing and Stimulation"

**The PDF file includes:**

Materials and Methods
Figs. S1 to S36
Tables S1 to S3
Supplementary Sections S1-S3
References 1-54

**Other Supplementary Materials for this manuscript include the following:**

Movies S1 to S4

### Materials and Methods

#### Materials

Four types of Tuftec™ styrene ethylene butylene styrene copolymer (SEBS) (H1062, H10521, H1052, H1221) were obtained from Asahi Kasei. Photoresist (AZ®1512) was obtained from MicroChemcials, UK. Microposit™ Developer MF-CD-26 was obtained from Micro Resist Technology GmbH, Germany. Polyimide solution (PI2611) was obtained from HD MicroSystem. Polyimide film (Kapton® Polyimide Film, Electrical-Grade, 12" Wide, 0.0050" Thick, 12" Long) was obtained from McMASTER-CARR. The following materials were all obtained from Sigma Aldrich: Poly(3,4-ethylenedioxythiophene)-poly(styrenesulfonate) (PEDOT PSS, product # 739332), SBS (product # 182877), poly(ethylene glycol)-block-poly(propylene glycol)-block-poly(ethylene glycol) diacrylate (PEG-PPG-PEG DA, product # 915858), lithium phenyl-2,4,6-trimethylbenzoylphosphinate (product # 900889), phenylbis(2,4,6-trimethylbenzoyl)phosphine oxide (BAPO, product # 511447), pentaerythritol tetrakis(3-mercaptopropionate), (PETMP, product # 381462), dimethyl sulfoxide (DMSO, product # 276855), nitric acid 25% (product # 1.60317), eutectic gallium–Indium (EGaIn) (product # 495425), AgNO3 (product #209139), KNO3 (product # 221295), FeCl3 (product # 157740), Polyvinyl butyral (PVB) (product # P110010), dextran (product # 31417), Nafion™ 117 containing solution (product # 70160), aniline (product # 242284), phosphate buffered saline (PBS) (product # 806552), and fluoxetine hydrochloride (SSRI) (product # F132).

#### Fabrication and characterization

Preparation and transformation of 2D films into fibers: A water solution of dextran was spin-coated (50 mg/ml, 1000 rpm, 1 min) on an oxygen plasma treated Si wafer and baked at 150 °C for 5 min. Then, a filtered toluene solution of SEBS was spin coated and followed by baking at 130 °C for 10 min. Different SEBS concentration (150 mg/ml, 100 mg/ml, and 50 mg/ml) were used to prepare 2D films of different thicknesses. Different types of SEBS (H1062, H10521, H1052, H1221) were used to create films of varying moduli. The 2D film was then released from the Si wafer by immersion in water for 1 hr and then transferred to an octadecyltrichlorosilane (OTS)-treated Si. The sample was then dried under 70 °C for 1 hr. The edges were cut using a blade. The transformation process then started by forming the core of the fiber using a glass slide to scrap against the edge of the film. Once a 30–50-micron core was achieved, we started the rolling process using a glass slide as a moving stage to obtain the final fiber. The fiber was annealed under vacuum (-70 cmHg) at 100 °C overnight.

To control the adhesion between the 2D film and the stationary substrate, OTS or oxygen plasma treated Si wafers were used. The rolling process could not properly proceed on the oxygen plasma treated Si as it led to 2D film tearing. To control the adhesion between the moving stage (glass) and the forming fiber, oxygen plasma treatment (1 min) was used. In addition, gentle pressure was applied during the rolling process to tighten the packing of the different layers of the fiber. High adhesion (between moving stage and the fiber core) and lower pressure led to fiber sliding and improper fiber rolling.

Preparation of light guiding and fluid delivery fibers: For light guiding fibers, a 2D film of SEBS (H1062) was prepared as previously described. A 250 µm diameter PMMA optical fiber (AZIMOM PMMA Plastic End Glow Fiber Optic Cable) was used as a core for starting the rolling process, skipping the scraping part of the transformation process. For fluid delivery fibers, a 2D film of SEBS (H1062) was prepared and cut. Then, a film of a home-made polypropylene glycol (PPG)-based polyurethane urea (PPG-PUU) (slightly modified from a previously reported polymer (*1*): 1:2:1 molar ratio of diamine-terminated poly(propylene glycol) (Jeffamine), isophorone diisocyanate (Thermo Scientific Chemical, product # 427600500), and 1,10-decanediol (Sigma Aldrich, product # D1203), respectively) was sprayed (from a solution of 20 mg/ml in isopropanol) next to the SEBS film, with a very small overlap area. The rolling process started from the PPG-PUU side, which ended up in the core of the fiber. The fiber was annealed under vacuum (-70 cmHg) at 100 °C for overnight, and then developed in acetone to dissolve the PPG-PUU core and obtain a hollow SEBS fiber.

Preparation of polyimide fibers: For polyimide_fiber preparation, a diluted polyimide_solution (1:1.5 diluted in NMP) was directly spin coated (5000 rpm, 1 min) on an OTS treated Si wafer and baked at gradually increasing temperature, 100 °C to 300 °C (25 ° / 10 min) under $N_2$. PPG-PUU (50 mg/ml in IPA) was then spin coated (2000 rpm, 1 min) as an adhesive layer and baked for 10 min at 120 °C. SEBS or PPG-PU were sprayed at the edge of the 2D film to be used for forming the fibrous core for rolling. This step was necessary since polyimide films could not be scraped into a fibrous core, as previously described with the elastic films.

Laser-induced graphene (LIG) sensor preparation and characterization: direct laser carbonization of a polyimide substrate was used to generate LIG patterns, which constitutes the main sensing components. Kapton films were laser engraved by an Epilog Fusion M2 $CO_2$ Laser following a reported method on LIG (*2*). A laser power of 7.5 W was used with a raster speed of 30 %. For morphological and compositional characterization of LIG, SEM images were taken by a FEI XL30 Sirion SEM (5 kV). The LabRAM HR Evolution microscope (HORIBA Scientific) was used for Raman spectroscopy with 532 or 633 nm excitation and a 1,800 grooves $mm^{-1}$ grating.

Mechanical property characterization: Mechanical properties of the bioelectronic fibers were studied using Instron 5565 with a 100-N loading cell. Fibers with different diameters (180, 270, and 360 µm) or made of different types of SEBS (H1062, H1052, H1051, H1221) were tested. The length of the fiber was measured by a caliper. The thickness was measured by an optical microscope. A strain rate of 100 mm/min was used for mechanical testing.

Electrical and mechanical property characterization of EGaIn: To measure the conductivity of the EGaIn electrodes during stretching, samples were attached to a homemade stretching station attached to an LCR meter (Keysight Technologies E4980). Silver epoxy was used to make contacts. The contact area was sealed with Torr Seal® epoxy to avoid strains on that area. EGaIn traces with a size of $0.5 \times 20\ mm^2$ were used to measure the resistance under stretching from 0 – 50 % in 5% steps.

Preparation of the GI S-NeuroStrings: Dextran (50 mg/ml) in water was spin-coated (1000 rpm, 1 min) on a Si wafer and baked under 150 °C for 5 min. Then, SEBS (H1062, 80 mg/ml) with 4% (w/w) azide-based cross-linker, bis(6-((4-azido-2,3,5,6-tetrafluorobenzoyl)oxy)hexyl) decanedioate,(*3*) in toluene was spin-coated (1000 rpm, 1 min). This was followed by 10 min baking under 150 °C. The desirable functional components (LIG and PEDOT-PSS sensing/stimulation interfaces and EGaIn interconnects) were then patterned on the SEBS film (described in detail in the next section). Finally, the patterning of the SEBS (H1062) encapsulation layer was done using spray coating (1 ml of 25 mg/ml SEBS solution) through a shadow mask. The film was annealed at 135 °C under vacuum (-70 cmHg) for 5 hr. The 2D sheet containing functional components was released from the Si wafer by immersion in water for 1 hr and then transferred to an OTS-treated Si. The sample was then dried under 70 °C for 1 hr. The edges were cut using a blade and then the film was manually rolled using a glass slide, as shown in Movie S1. The connection area was kept unrolled for subsequent connection with a flexible flat cable. After connection and sealing, the fiber was subjected to further modifications such as silver and PANI electrodeposition, immersion in $FeCl_3$, and coating with PVB and Nafion (fully described in the next sections) for obtaining the desirable sensing components. The fibers were annealed under vacuum (-70 cmHg) at 100 °C before use. This fabrication process provided a yield of > 95%, as estimated by the percentage of working functional components.

<u>EGaIn interconnects patterning:</u> Photoresist (AZ®1512) was spin-coated (2500 rpm for 45 sec) on a spin-coated SEBS (H1062) substrate and followed by light exposure (385 nm, 90 mJ/cm$^2$) using Durham Magneto Optics ML3 MicroWriter. After light exposure, the photoresist was developed for 45 sec in Microposit™ Developer MF-CD-26. Then, gold (30 nm) was thermally evaporated on the surface to provide an adhesion layer between EGaIn and SEBS. EGaIn was screen-printed onto the surface using a PDMS as the blade. This process is repeated multiple times until a smooth EGaIn film was obtained. The sample was immersed in acetone for 30 min and followed by 30 sec of sonication. This was repeated until the photoresist layer was completely removed.

<u>Preparation and characterization of pressure/stain sensors:</u> LIG was transferred to an SEBS (H1062) film by laminating a PI film with LIG on it. Pressure was applied using a metal spatula on the PI film to achieve better transferring of LIG. Then, EGaIn interconnects were patterned to form resistors for use as piezoresistive pressure/strain sensors and followed by spray coated SEBS (H1062) encapsulation. After film release and rolling, the sensors were in the desired position inside the fiber according to the design layout and rolling direction. The performance of these sensors after rolling was examined using different tests: i) normal force (0.1 N – 2 N) was applied directly on the sensor using a force gauge. ii). A metal cylinder was manually rolled on the fiber at different speeds to create progressing forces. iii) In-tube force sensing demonstration: the fiber was inserted into a Tygon tube (Tygon® E-3603; wall thickness: 0.062 in; external diameter: 0.5 in, modulus: 12.1 MPa). External force was applied on the wall of the tube to cause structural deformation that mimics the conditions in the intestine. The resistance was measured and correlated with the applied forces. In all tests, the current (at 0.5 V) was recorded using PalmSens4 (PalmSens BV, Netherlands). For signal processing, a baseline was first generated using the asymmetric least squares smoothing function in OriginLab. Then, the signal was normalized with the baseline: $\frac{\Delta I(t)}{I_0(t)}\% = \frac{I(t) - I_0(t)}{I_0(t)} \cdot 100$, where $I$ is the raw current signal and $I_0$ is the calculated baseline.

<u>Preparation and characterization of serotonin sensors:</u> A two-electrode electrochemical configuration was used for serotonin sensing. The working electrode was based on LIG (0.5 mm$^2$) embedded on the surface of SEBS (prepared by transferring from polyimide). Nafion (5% in a mixture of lower aliphatic alcohols and water) was electrodeposited onto the serotonin sensing electrodes to minimize the effect of interfering molecules (e.g. ascorbic acid) according to a previously reported method.(*4*) Briefly, the electrodes were immersed in Nafion solution under a constant voltage (1V) against an Ag/AgCl commercial reference electrode (Sigma Aldrich,

product # BASMF2056) for 30 seconds.  The electrodes were washed with water and dried at 100 $^{o}$C for 10 min. For the reference electrode (located at the tip after fiber rolling), Ag/AgCl based electrode was prepared by silver electrodeposition on LIG (5 $mm^2$) from an aqueous solution of 5 mM AgNO3 and 1 M KNO3. The LIG surface electrode was dipped into the electrodeposition solution. The potential was then swept from −0.9 to 0.9 V versus an Ag wire (0.5 mm in diameter) for 20 segments at a scan rate of 0.1 V $s^{-1}$.(*5*) After washing with water, the surface of silver was immersed in FeCl3 solution (3M) for 10 sec to form the AgCl. Finally, the electrode was covered with a PVB layer for stabilization by dipping in a methanol solution of PVB (50 mg/ml) (*6*-*8*). After transforming the 2D film into fiber, on-fiber serotonin sensors were characterized *in-vitro* using chronoamperometry (at 0.5 V) or cyclic voltammetry (-0.4 to 1.3 V, 10 V/sec, 0.001 V step) under different concentrations: 100, 200, 400, and 800 nM.

<u>Preparation of and characterization pH sensors:</u> For pH sensing, a potentiometric setup was used with a two-electrode cell. The working electrode was based on polyaniline (PANI)-modified LIG. PANI was electrodeposited using a previously reported method.(*9*) The same previously described reference electrode was used also here. The pH dependence was measured as PANI known to be sensitive to pH due to different degree of doping (*10*). This intrinsic sensitivity was used to translate pH values to changes in open-circuit potential (OCP). The pH sensor was characterized in buffer solutions with varying pH values ranging from 3-10. The OCP was measured for 30 sec in each solution. The measurement was paused while transferring the electrodes from one solution to another.

<u>Preparation and characterization of PEDOT-PSS-based stimulation electrodes:</u> A solution of PEDOT-PSS (1.1% in H2O), PEG-PPG-PEG DA (Mn ~5,800), and lithium phenyl-2,4,6-trimethylbenzoylphosphinate (100 : 2 : 0.05 by weight, respectively) was prepared and stored in the fridge. 10 µl of flurosurfactant (Capstone™ FS-30) (for better wettability) and 50 µl of DMSO (for better conductivity) were added to each 1g of solution before use. The final mixture was sprayed (0.5 ml) on the surface through a laser-cut shadow mask (with openings of 5 × 3 $mm^2$) to obtain the stimulation interface. The film was cured under UV (250 W iron-doped lamp, 365 nm, Honle UV America Inc.) under $N_2$ for 20 min and then baked for another 10 min under 135 °C. The film was immersed in methanol for 1 min and then baked at 135 °C for 2 min. The film was then immersed in nitric acid (25%) for 30 sec, washed with water, and dried on a hot plate (135 °C) for 1 min. EGaIn was then patterned to form the interconnects and followed by encapsulation with an SEBS (H1062) layer by spray coating. For characterization, two PEDOT-PSS based stimulation electrodes were immersed in PBS. Impedance was measured from 1MHz to 1Hz. Stimulation currents were also measured under biphasic stimulation conditions: 5 ms pulse duration at different 0.5, 1, 2, and 4 V.

<u>Preparation of brain S-NeuroString:</u> Dextran (50 mg/ml) in water was spin-coated (1000 rpm, 1 min) on a Si wafer and baked under 150 °C for 5 min. Then, SBS (80 mg/ml) with 4 wt% PETMP, and 4 wt% BAPO (vs. SBS weight) (*3*), in toluene was spin-coated (1000 rpm, 1 min). The film was cured under UV (250 W iron-doped lamp, 365 nm, Honle UV America Inc.) for 1 min. This was followed by 10 min baking under 135 °C. The film was cooled down and immersed in toluene for 3 min to remove any unreacted residues. This was followed by 2 min baking at 135 °C and oxygen plasma treatment. PEDOT-PSS based electrodes were then patterned. The previously described PEDOT-PSS solution (PEDOT-PSS (1.1% in H2O), PEG-PPG-PEG DA (Mn ~5,800), and lithium phenyl-2,4,6-trimethylbenzoylphosphinate (100 : 1 : 0.05 by weight, respectively)) was spin coated (1000 rpm, 90 sec) on the plasma-treated SBS film. The film was cured under UV

(250 W iron-doped lamp, 365 nm, Honle UV America Inc.) under $N_2$ for 20 min and then baked for another 10 min under 135 °C. The film was immersed in methanol for 1 min and then baked at 135 °C for 2 min. A very thin layer of SEBS (H1062) was spin coated (10 mg/ml in toluene, 2500 rpm, and 1 min) to achieve a better adhesion between the PEDOT and the subsequent photoresist layer. After baking (135 °C, 1 min), a photoresist layer was spin coated (2500 rpm, 45 sec), followed by light exposure (385 nm, 90 mJ/cm$^2$) using Durham Magneto Optics ML3 MicroWriter at a writing speed of 180 mm$^2$/min, and then development for 45 sec in Microposit™ Developer MF-CD-26. The exposed PEDOT layer was etched using plasma etching (March Instruments PX-250 Plasma Asher, 150 W, 250 sec). The film was immersed in acetone for 30 seconds under sonication to remove the photoresist layer and followed by development with cyclohexane for removing the remaining SEBS. The PEDOT-PSS electrodes were then immersed in nitric acid (25% by weight) for 30 sec, washed with water, and dried on a hot plate (135 °C) for 1 min. SBS was then directly photopatterned for encapsulation. Briefly, SBS solution (80 mg/ml in toluene) was spin coated and exposed using ML3 MicroWriter (385 nm, 350 mJ/cm$^2$) at a writing speed of 180 mm$^2$/min. The film was baked for 2 min at 90 °C and then developed using cyclohexane under spinning (2500 rpm, 45 sec). This step was repeated to give a double layer of SBS encapsulation with a total thickness of 3.2 micron. The film was baked at 135 °C for 10 min. The 2D sheet containing all recording electrodes was released from the Si wafer by immersion in water for 1 hr and transferred to an OTS-treated Si. The sample was then dried under 70 °C for 1 hr. The edges were cut using a laser cutter. After cutting, the film was manually rolled using a glass slide. The film has a varying width (with the smallest width at the recording area), which led to a fiber with a thinner tip. The connection area was kept unrolled for subsequent connection with a flexible flat cable. After connection and sealing, the fibers were annealed under vacuum (-70 cmHg) at 100 °C for overnight before use. This fabrication process gave a yield of > 90% for each batch, as estimated by the percentage of recording electrodes with an impedance value of < 1MΩ at 1kHz.

Impedance tests: For long term size-dependent impedance test, 4 sizes of PEDOT electrodes were prepared (25 µm x 25 µm, 50 µm x 50 µm, 100 µm x 100 µm, and 200 µm x 200 µm). Impedance was measured versus a commercial platinum wire (0.5 mm in diameter) from 1MHz to 1Hz. This test was done at different time points after immersion (immediately, 1 week, 2 weeks, and 1 month). For the encapsulation test, fully encapsulated (a double layer 3.2-micron-thick SBS) PEDOT-PSS based electrodes were used. Impedance was measured versus a commercial platinum wire from 1MHz to 1Hz to show the robustness of the encapsulation. This test was also done at different time points (immediately, 1 week, 2 weeks, and 1 month). For the fiber microelectrodes impedance test, the impedance of the 32 on-fiber microelectrodes were measured in PBS solution versus a platinum wire electrode from 1MHz to 1Hz.

**Gastrointestinal Studies**

All surgical procedures for mice and pigs were performed in accordance with protocols (32497, 32778, 34283, and 31893) approved by the Institutional Animal Care and Use Committee (IACUC) at Stanford University.

*Ex-vivo* study of motility and stimulation: Spatiotemporal mapping was completed according to a published protocol.(*11*) Mice were fasted overnight and euthanized by carbon dioxide administration and cervical dislocation. Colons were removed and pinned loosely in a bath containing 37°C physiological Krebs solution. A camera (DMK 41AF02, The Imaging Source,

Charlotte, North Carolina) was oriented above the colon and used to record 5-min videos (3.75 fps, 1280x960, 8-bit) using an IC Capture software (The Imaging Source, version 2.4.642.2631). A bifunctional S-NeuroString (300 µm in diameter and 5 MPa Young's modulus) with 8 internal pressure (0.5 cm distance between sensors) sensors and 4 surface stimulation electrodes (1 cm distance between electrodes) was inserted into the colon for motility sensing and stimulation. On each colon (n=5 mice), we recorded under three different conditions (5 min each): 1) empty colon, 2) colon with S-NeuroString, 3) Colon with S-NeuroString under Stimulation. We used a biphasic stimulation (2V vs AgCl ref electrode, stimulation pulse: 5 msec, stimulation period: 5 sec) to trigger colonic motility by all stimulation channels simultaneously. Five stimulation cycles were done (at t = 30, 90, 150, 210, 270 sec) during the 5 min recording period. Simultaneously, the change in colonic diameter was captured by the camera and pressure sensing was done using the S-NeuroString.

The output signals obtained from the pressure sensors were processed as described in the previous sections. For motility peak detection (peak location, amplitude, duration, and area under the peak), we used the peak analyzer in OriginLab. Video analysis was performed using a custom-written software (VolumetryG9a, Dr. Grant Hennig, University of Vermont). In brief, videos of the colons were transformed into particle formats, and a spline was fit longitudinally to measure the diameter over time at every point along the length of the colon. Spatiotemporal maps, which display the matrix of diameter versus time, were generated. From these maps, two thresholds were set to select for long (83 frames, 8 to 30 seconds) and short (9 frames, 0.8 to 3.2 seconds) intervals between contractions to identify (colonic migrating motor complexes) CMMCs and slow wave contraction, respectively. A color gradient was fit to the spatiotemporal maps to represent the interval between slow waves at each cross-sectional diameter for the entire length of the colon. The summation of each focal diameter over the 10-min time period was normalized to graph the cumulative distribution of the percentile of intervals (in seconds) of CMMCs and slow wave contractions. The procedure was followed for recording motility signals in the intestine. For the effect of the fiber on colonic motility, the fiber was inserted half-way into the colon. The camera was used for motility recording. The above procedures were used for fig. S22, fig. S24, and fig. S25.

*In-vivo* colonic motility recording in mice: Six- to eight-week-old C57BL/6 mice obtained from Charles River Labs (Wilmington, MA) were used in this study. Mice (n=5) were anesthetized with 2-3% inhaled isoflurane. The colons of anesthetized mice were washed using a sterile saline solution. S-NeuroString (with 6 pressure sensors and 0.5 cm distance between sensors) was then inserted into the colon, lubricated by the sterile saline wash, via the anus (3-4 cm deep). Since S-Neurostring is slightly stiffer than the colonic tissue but still relatively soft, it was easy to push the device into the colon without causing damage (fig. S26 B-D). Histology confirmed that no damage was caused during the insertion of the fiber (fig. S23). Motility was recorded for 10 min. The mice were then euthanized and colons were dissected to verify the location of S-NeuroString and to make sure the colonic tissue was not perforated. The above procedures were used for fig. S26.

*In-vivo* motility sensing and stimulation in anesthetized pigs: Female juvenile Yucatan miniature pigs (Sus scrofa) aged 6–9 weeks and weighing 10 kg from breeder S&S Farms (Ramona, CA) were used in this study. All pigs were fasted overnight, placed under general anesthesia using vaporized isoflurane and connected to vital-sign monitors during this terminal procedure. A midline laparotomy incision was made to expose the viscera. Krebs/PBS buffer was infused into the porcine intestine to flush the contents before sensing. The ligament of Treitz was identified to locate the duodenum and jejunum and the ileocecal valve was found to designate the ileum. For recording natural motility, a soft 10-cm long S-NeuroString holding 8 pressure sensors was inserted within the lumen of the small intestine through an ostomy, for continuous live recordings. Natural motility was infrequent and unpredictable due to the effect of anesthesia, but we managed to record 3 times. For studying electrical stimulation, a bifunctional S-NeuroString with (5 stimulation electrodes and 5 pressure sensors) was used. Baseline motility was recorded for 30 second followed by biphasic stimulation at 2V and 4V (stimulation pulse 5 msec, stimulation period: 5 sec). This experiment was done on 2 pigs on multiple intestinal location (n = 7 stimulation experiments). For the programable stimulation, pressure sensing was done using the distributed sensors (S1-S5). Selective stimulation was done by activating a single stimulation electrode each time, starting from SE1, SE3, and then SE5. The above procedures were used for Fig. 3C, F, G, I, and fig. S27.

<u>Electrochemical sensing of serotonin in anesthetized pigs:</u> Female juvenile Yucatan miniature pigs (Sus scrofa) aged 6–9 weeks and weighing 10 kg from breeder S&S Farms (Ramona, CA) were used in this study. Briefly, all pigs were fasted overnight, placed under general anesthesia using vaporized isoflurane and connected to vital-sign monitors during this terminal procedure. A midline laparotomy incision was made to expose the viscera. A PBS solution was infused into the porcine intestine to flush the contents. The ligament of Treitz was identified to locate the duodenum and jejunum and the ileocecal valve was found to designate the ileum. A soft 10-cm long S-NeuroString with 5 serotonin sensors was inserted within the lumen of the small intestine through ostomies, for continuous recordings. Stabilization of the baseline was then performed for 10 min after the injection of PBS (0.5 ml). After 10 min, a new baseline was recorded for 1 min. SSRI (fluoxetine hydrochloride) solution (10 µM, 0.5 ml) was then administered upstream to the sensing area by direct injection into the lumen. Measurements were performed 10 min after the injection. This experiment was done on 3 pigs on multiple intestinal segments (n = 5). Anaesthetized animals were euthanized by trained researchers at the end of the recordings. The data was collected from all 5 sensors simultaneously. However, not all sensors were responsive, i.e., some sensors have shown no change after SSRI injection while others responded strongly (fig. S28). We assume that this may be related to the distance of the fiber from the wall of the intestine or its orientation, which was not well-controlled in this experiment and is known to affect the sensitivity to serotonin (*12*, *13*). A single sensor may not capture all changes in concentrations accurately due to its localized measurement. However, by using multiple sensors, we can take advantage of their distribution and obtain a more reliable average concentration based on multiple readings as shown in Fig. 4K.

To verify that the signal is coming from serotonin, we used cyclic voltammetry. The fiber was inserted into the intestine as described earlier. Stabilization of the baseline was then performed by continuous cyclic voltammetry (V: -0.4 to 1.3V, scan rate: 10 V/sec) for 10 min after the injection of PBS (0.5 ml). After 10 min, a new baseline was recorded which defined the background current in the intestine. SSRI solution (10 µM, 0.5 ml) was then administered upstream to the sensing area by direct injection into the lumen. Measurements were performed immediately and 10 min after injection, showing the appearance of serotonin peaks. To further prove that the peaks originate from serotonin, we injected a solution of serotonin hydrochloride (10 µM, 0.5 ml) and showed that the peaks (obtain after SSRI injection) became larger. The above procedures were used for Fig. 3K,L, and fig. S28.

<u>Histology and biocompatibility studies:</u> Three groups (n=5) of six- to eight-week-old C57BL/6 mice obtained from Charles River labs were used in this test. Mice were anesthetized via the

inhalation of 2-3% isoflurane. The colons of these mice were washed using sterile saline. This was followed by the insertion of the S-NeuroString (length 2 cm, diameter 300 µm) or a flexible polyimide film (Kapton® from McMASTER-CARR, length: 2 cm, width: 300 µm, and thickness: 127 µm). Another group of 5 mice (sham) was used as a control; in these mice, no probe was inserted after washing the colon. During the insertion process, the probe was fully inserted and fully removed 10 times from the colon. Horizontal and vertical movements (10 times each) of the device were performed to simulate ambulation. Once the probes were fully inserted, force was applied on the abdominal area by applying manual pressure 10 times to simulate GI contractions. The mice were then awakened for 30 mins, during which time their free movements were tracked and the number of passed stool pellets were counted. The mice were immediately euthanized following the study. The colons were then collected, and the tissue was fixed in neutral buffer with 10% formalin (vol./vol.) for 15 h, transferred into 70% ethanol for 5 h and embedded in paraffin. The tissue sections with thickness of 5 µm were prepared and stained with H&E and trichrome staining (completed by the Stanford Department of Comparative Medicine's Animal Histology Services). The stained sections were scanned by a Hamamatsu NanoZoomer 2.0-HT and the images were processed using ImageScope viewing software. The above procedures were used for fig. S23.

Locomotor and pellet output behavior: Following the insertion process described in the previous section, where the bioelectronic fiber was placed in the colon of one group, flexible Kapton was placed in the colon of the second group and no device was placed in the colon of the third group, the locomotor and pellet output behavior of the mice were characterized. After recovering from anesthesia, mice were placed in a chamber, in which they freely moved. Their open-field activity was assessed on the basis of the below cited previously published protocol (*14*). The mice were videotaped for 30 min using a smart phone camera and the movement was analyzed by MATLAB. The number of stool pellets passed by the mice were counted manually. The average residence time of fibers in the colon was also assessed. The above procedures were used for fig. S23.

**Brain studies**

Implantation of neural S-NeuroString: All experiments performed on mice were approved by Stanford University's Administrative Panel on Laboratory Animal Care (34010). The animal care and use programs at Stanford University meet the requirements of all federal and state regulations governing the humane care and use of laboratory animals including the United States Department of Agriculture Animal Welfare Act and the Public Health Service Policy on Humane Care and Use of Laboratory Animals. The laboratory animal care program at Stanford is accredited by the Association for the Assessment and Accreditation of Laboratory Animal Care. All mice were maintained on a reverse 12-hour dark/12-hour light cycle (temperature: 20° to 25°C; humidity: 50 to 65%) in the Stanford University's Veterinary Service Center and fed with food and water ad libitum as appropriate. All experiments occurred during their active cycle.

Anesthesia was induced with isoflurane (4%; maintained at 1.5%). Once anesthetized, the mouse was head-fixed on a stereotactic frame (RWD) and body temperature was maintained with a heating pad (Keenovo). The eyes were protected with ophthalmic ointment (Puralube). The skull was exposed, and a 0.5-mm-diameter craniotomy was made at the following coordinates from bregma: anteroposterior, -1.7 mm; mediolateral, 1.6 mm for the hippocampus (for recording and histology), and anteroposterior, 2 mm; mediolateral, 0.5 mm for the prefrontal cortex (for recording). A durotomy was then performed. The fiber device was then inserted at a velocity of 0.01 mm/s with simultaneous ultrasonic vibration (NeuralGlider) to the target depth of 3 mm for both the hippocampus and prefrontal cortex, and the cranial window was subsequently filled with Kwik-Sil (WPI). For electrical recordings, a self-tapping bone screw (Fine Science Tools) was secured over the cerebellum as the ground electrode, and a stainless steel headbar was cemented onto the skull using dental cement (C&B Metabond). After implantation, mice were habituated to run on a treadmill for 7 days. For the histology study, the fiber was fixed to the skull using dental cement and then the skin was sutured.

Electrophysiological recordings: The mice were placed on the treadmill and fixed to a headbar clamp. The probe was then connected to the data acquisition system (Intan RHS Stim/Recording System) through a custom adapter board. Electrophysiological data was collected at 30 kHz for 10 minutes._Data analysis was performed using custom software written in MATLAB 2019b (MathWorks). Spike sorting was performed using Kilosort 3.0 (https://github.com/MouseLand/Kilosort) with default parameters (*15*). Subsequently, quality metrics were computed and double counted spikes (within ±0.16 ms) removed using the ecephys spike sorting pipeline (https://github.com/AllenInstitute/ecephys_spike_sorting). Clusters were then manually curated in Phy (https://github.com/cortex-lab/phy). Single units were only classified as good if the ISI violations were less than 0.3 (*16*), the number of spikes were greater than 200, and the presence ratio was greater than 0.9. Signal-to-noise ratios were calculated as the amplitude of the spike on a single channel over the root mean squared value of the band-passed Butterworth filtered trace (400 to 7000 Hz). Dimensionality reduction was carried out by (i) collapsing each unit's waveform across all 32 channels into a single feature vector, (ii) principal component analysis to reduce it to 30 components, (iii) UMAP with the hyperparameters n_neighbors = 5, min_dist = 0.01, and spread = 1 (*17*).For sharp wave ripple detection, the electrophysiological data was band-pass Butterworth filtered (0.1 – 300 Hz), and downsampled to 1000 Hz for analysis. The raw LFP data was band-pass Butterworth filtered (125 – 200 Hz), and SWRs were classified if the envelope of the trace exceeded 4 SD for at least 15 ms (*18*).

Brain histology: Mice were anesthetized using pentobarbital sodium and phenytoin sodium (1 ul/1 g weight). Transcardial perfusion was performed with around 25 ml of 1×PBS, followed by 25 ml of 4% paraformaldehyde (PFA) one week after implantation. Subsequent to decapitation, the head was immersed in a 4% PFA solution for 24 hours. The brain was then removed from the skull, placed in a 4% PFA solution for another 24 hours, and subsequently transferred to a 30% (w/v) sucrose solution. The brain remained in the sucrose solution until it settled at the base of a 50 ml tube, ensuring complete dehydration. To safeguard the brain tissue for cutting, it was cryoprotected using the optimal cutting temperature (OCT) compound (Tissue-Tek, USA). Horizontal brain sections with 8 μm in thickness, were produced using a cryostat machine (Leica CM3050 S, Germany), and these sections were mounted onto glass slides.

Following cryostat sectioning, the brain slices underwent a 20-minute incubation in a solution containing 0.3% (v/v) Triton X-100 in PBST (0.1% Tween-20 in 1× PBS). After a PBST rinse, the slices were blocked with 5% (w/v) bovine serum albumin (BSA) at room temperature for a duration of 2 hours. Subsequent to PBST washing, the slices were subjected to an overnight incubation at 4 °C with primary antibodies targeting astrocytes (Rat anti-glial fibrillary protein GFAP, 1:1000, Abcam, ab279291), microglia (Rabbit anti-ionized calcium binding adaptor molecule 1 Iba1, 1:1000, Abcam, ab178846), and neurons (Chicken anti-neuronal nuclear NeuN, 1:1000, Sigma, ABN91). Following another PBST wash, the slices were exposed to secondary antibodies (1:1000, Alexa Fluor 647 goat anti-rabbit, Abcam, ab150087; 1:1000, Alexa Fluor 594 donkey anti-chicken, Jackson ImmunoResearch, 703-585-155; 1:1000, Alexa Fluor 488 goat anti-rat, Abcam, ab150165) in darkness at room temperature for a 2-hour duration. This incubation was followed by a PBST solution rinse. To preserve the samples, brain slices were sealed with an antifading mounting medium containing 4',6-diamidino-2-phenylindole (DAPI), covered with slips, and stored at -20 °C, shielded from light.

Imaging and Analysis of Image Data: Confocal fluorescence microscopy images were captured using Stellaris 8 DIVE upright confocal microscopes (Leica, Germany). For the antibody-labeled horizontal slices shown in fig. S35, confocal images were obtained using excitation sources with wavelengths of 405 nm, 488 nm, 561 nm, and 638 nm.

Statistical analysis: For data analysis, Origin, MATLAB, and Excel were used. All replicate numbers, error bars, P values and statistical tests were indicated in the figure legends.

**Supplementary Figures**

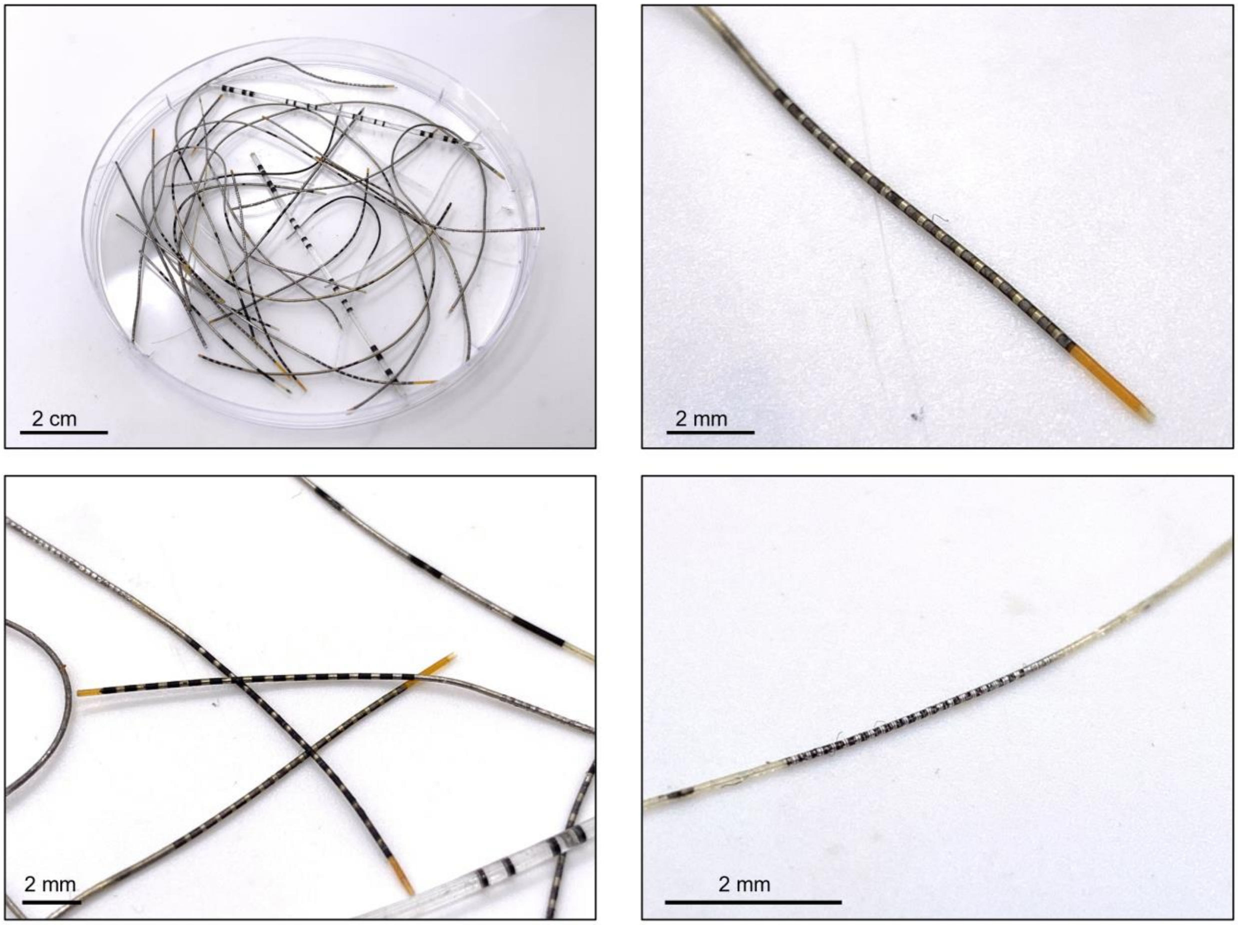


**Fig. S1**. Photographs of transformed electronic fibers with different sizes and configurations.

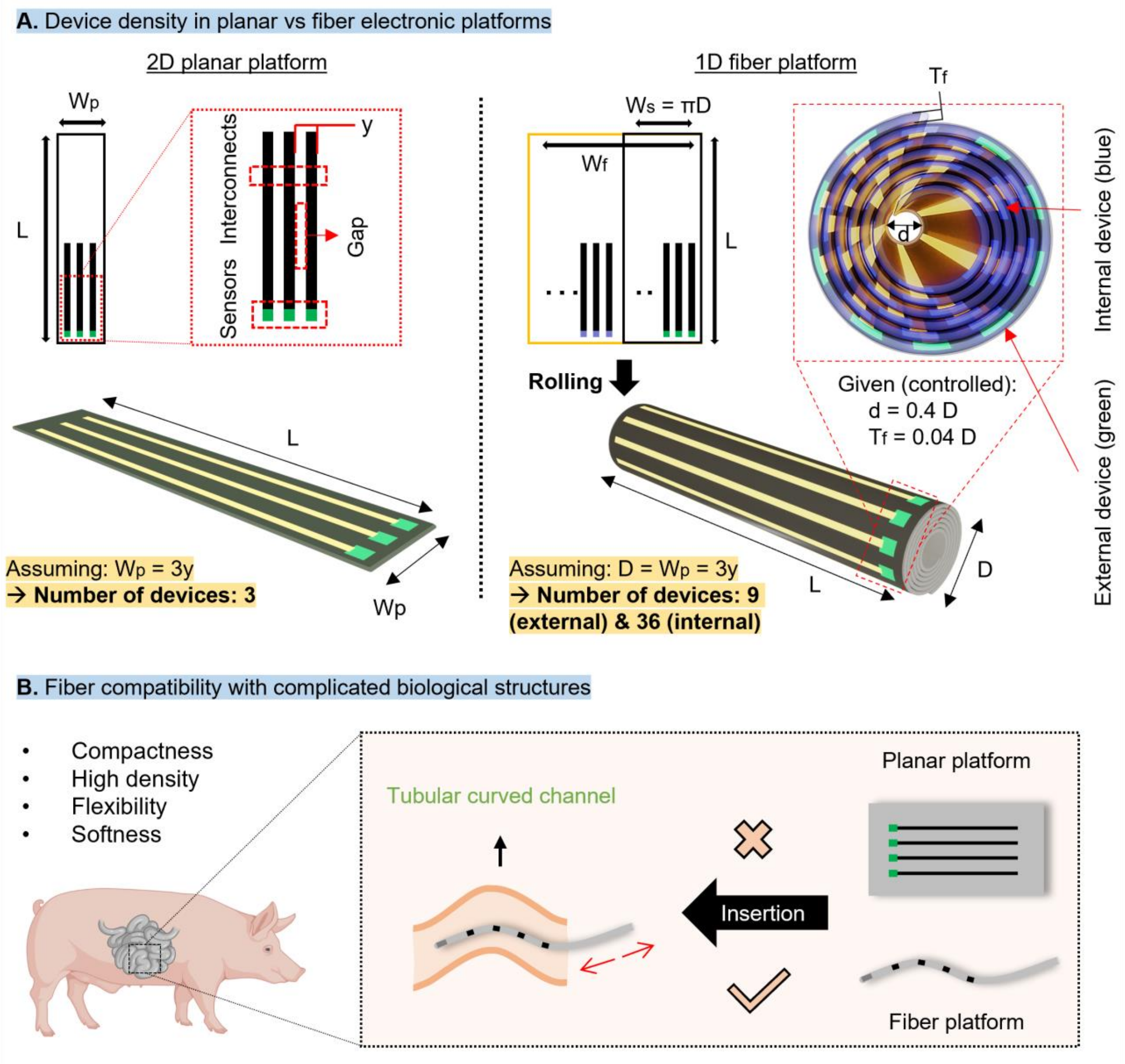


**Fig. S2. The rationale behind the development of bioelectronic fibers**. (**A**) A comparison between a 2D planar sheet and a 1D fiber platform. Assuming that the planar sheet and fiber have the same characteristic width (i.e., $W_p = D$) in this specific example (where the fabrication pitch is $y = 1/3\ W_p = 1/3\ D$, $d = 0.4\ D$, and $T_f = 0.04\ D$), the planar platform can host 3 components while the fiber hosts 9 surface (green) and 36 internal components (blue). $T_f$: thickness of the 2D (unrolled) film and d: internal diameter. (**B**) The advantages of fiber platforms in terms of compatibility with extended and torturous biological targets.

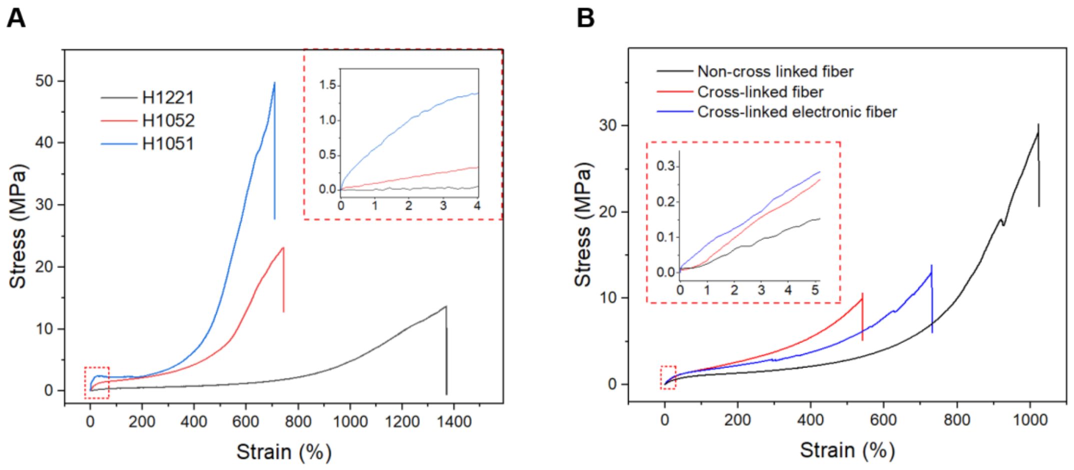


**Fig. S3. Mechanical properties of the S-NeuroStrings (300 μm diameter).** (**A**) Stress strain curves of fibers prepared from 3 different types of Tuftec™ SEBS (H1221, H1052, and H1051). (**B**) Stress-strain behavior of a cross-linked S-NeuroStrings (with 16 pressure sensors and EGaIn interconnects), compared to non-cross linked pure SEBS fiber. All fibers had the same diameter (350 μm).

| Sample | Maximum strain (%) | Stress at break (MPa) | Modulus (MPa) |
|---|---|---|---|
| SEBS (H1221) fiber | 1385 ± 15 | 14.0 ± 2.0 | 1.2 ± 0.2 |
| SEBS (H1052) fiber | 743 ± 68 | 20 ± 5.5 | 7.7 ± 0.9 |
| SEBS (H1051) fiber | 719 ± 96 | 53.3 ± 3.1 | 49.3 ± 6.4 |
| SEBS (H1062) fiber | 1027 ± 22 | 26.6 ± 2.6 | 2.6 ± 0.5 |
| Cross-linked (cl-) SEBS (H1062) fiber | 413 ± 26 | 7.9 ± 1.2 | 4.8 ± 0.6 |
| cl-SEBS (H1062) electronic fiber | 721 ± 254 | 13.4 ± 9.6 | 4.9 ± 0.3 |

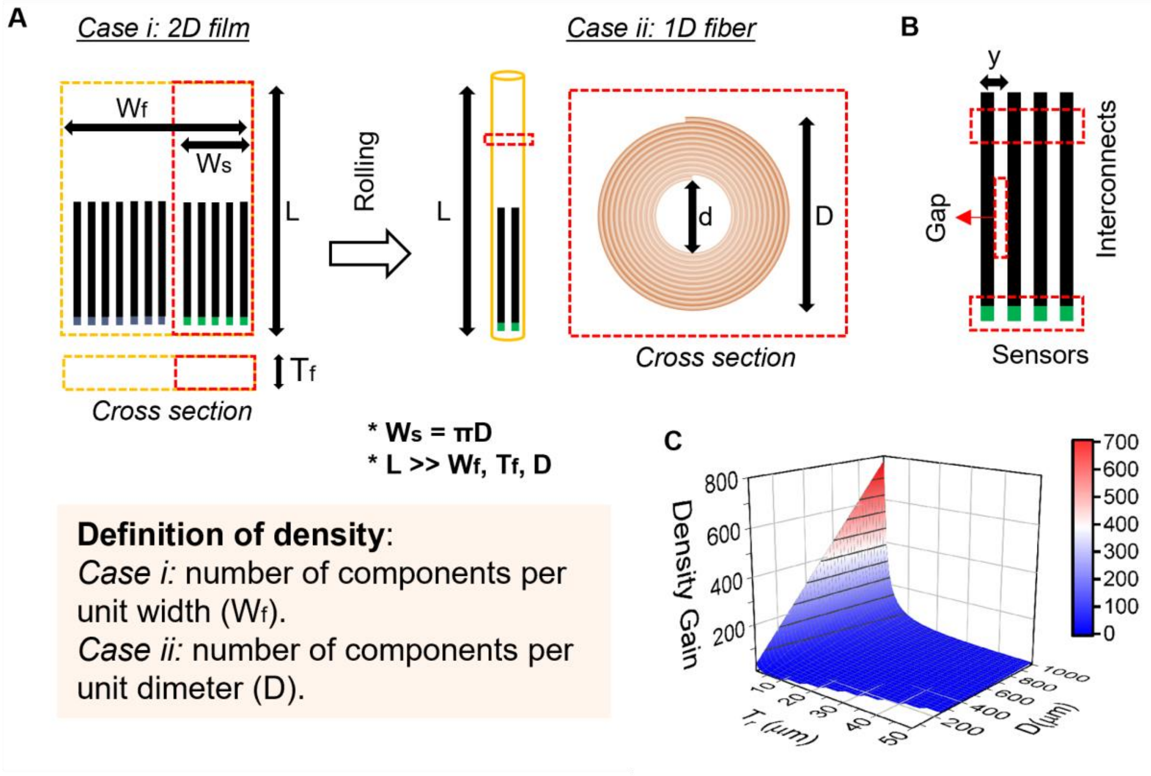


**Fig. S4. Density gain: A 2D-to-1D transformed fiber vs the corresponding 2D film.** (**A**) Scheme showing the 2D film before and after transformation into fiber. (**B**) Electronic component design and dimensions used for the calculation of density gain. Green squares represent sensors and black lines represent interconnects. The pitch size is denoted by "y". (**D**) Color map showing the density gain as a function of the unrolled 2D film thickness (Tf) and final fiber diameter (D).

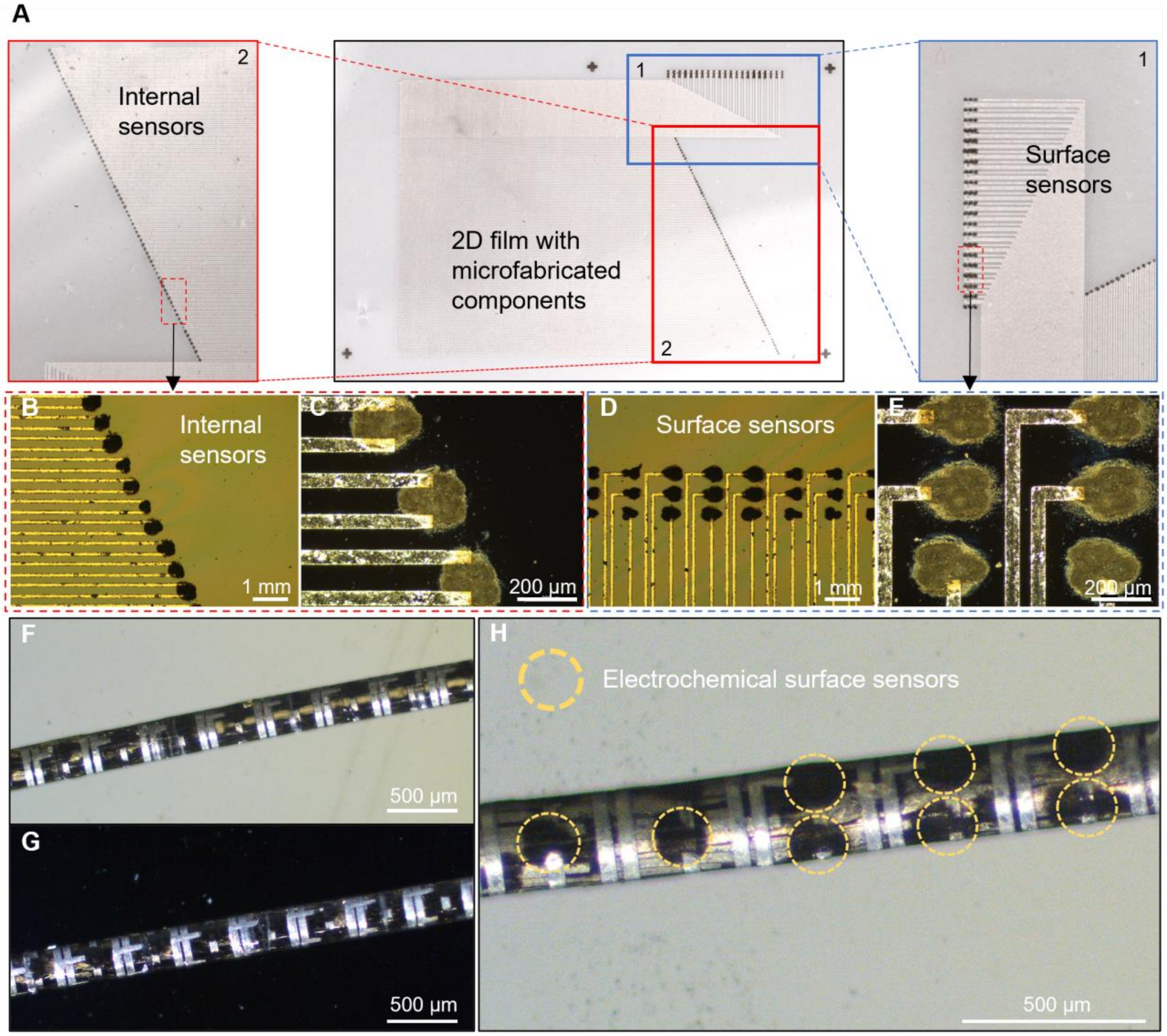


**Fig. S5. High density S-NeuroStrings.** (**A**) Photographs showing the layout of the 2D film before rolling into a high density fiber. (**B**)-(**E**) Microscope images of microfabricated LIG-based sensors (81 pressure and 69 electrochemical sensors) and EGaIn interconnects. (**F**)-(**H**) Images of the transformed fiber showing multiple internal and surface electrical components.

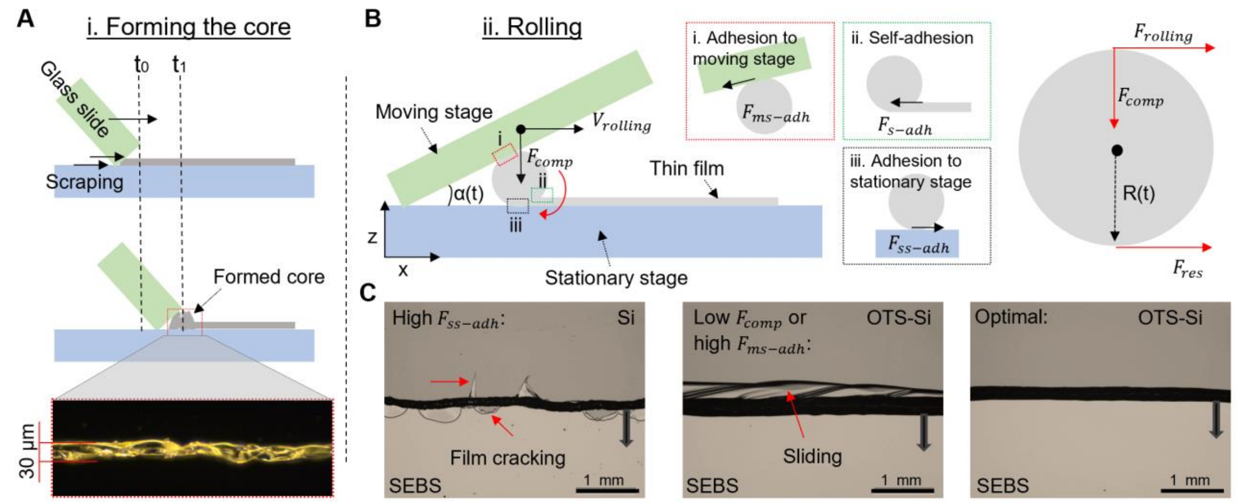


**Fig. S6. 2D-to-1D Transformed Fibers**. (**A**) Schematic illustration showing the first part of the transformation process where a 2D film is scraped to form a core which can be later used for rolling. (**B**) Schematic illustration showing the rolling process and the forces involved in the process. $F_{adh-ms}$ is the adhesion between the film and the moving substrate. $F_{adh-ss}$ is the adhesion between the film and the stationary substrate. $F_{adh-self}$ is the self-adhesion of SEBS (or the other used elastomers). $F_{rolling}$ is the force applied on the fiber to maintain the rolling procedure, $F_{res}$ represents forces that resist the rolling process (e.g. friction with the stationary substrate, rolling resistance due to fiber deformation, etc.), and $F_{comp}$, compressive force generated by the moving stage to help maintain good adhesion between the layers and prevent sliding. (**C**) Optimization of the rolling process by tuning different parameters such as $F_{adh-ss}$, $F_{adh-ms}$, and $F_{comp}$.

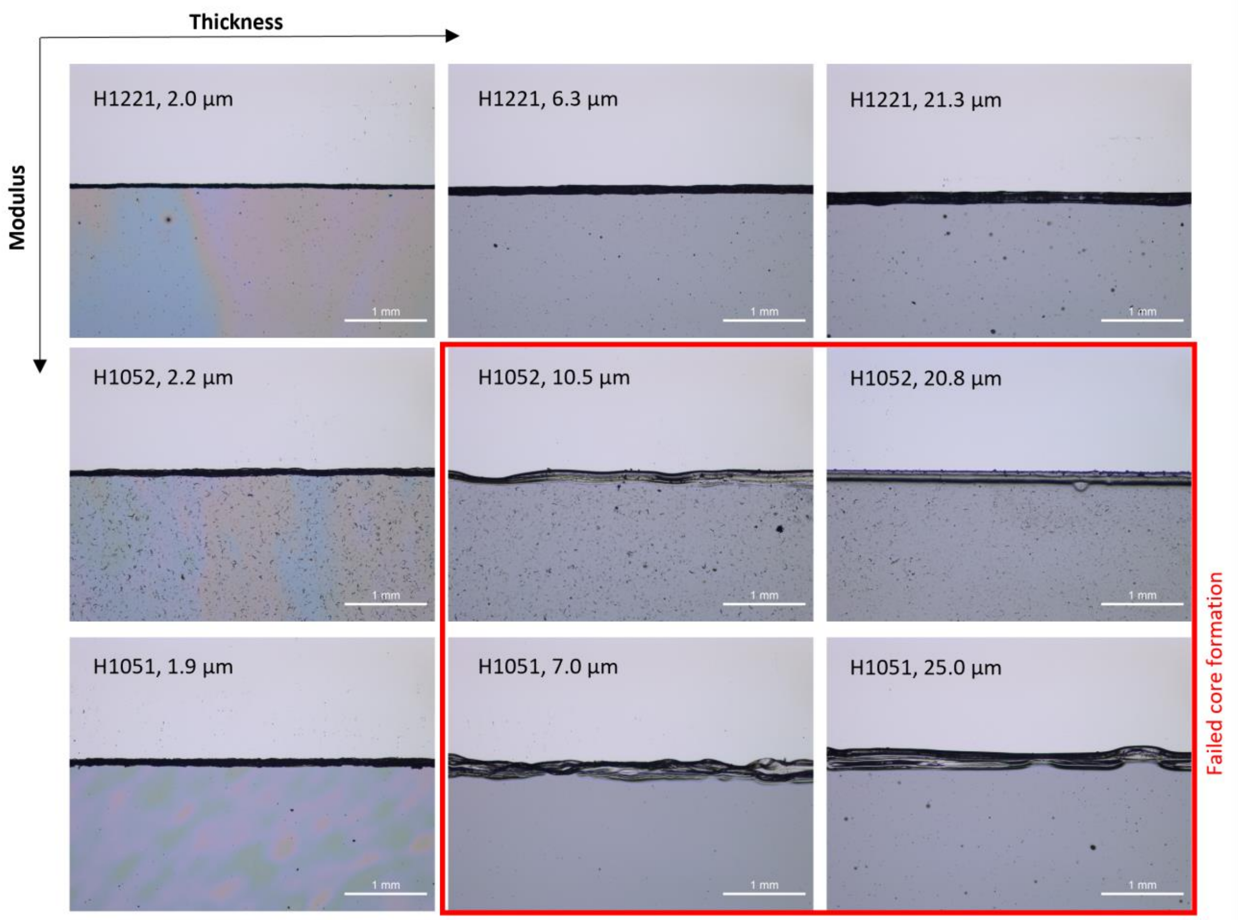


**Fig. S7. The effect of film modulus and thickness on the core formation process.** Thinner films and lower modulus allow for easier core formation.

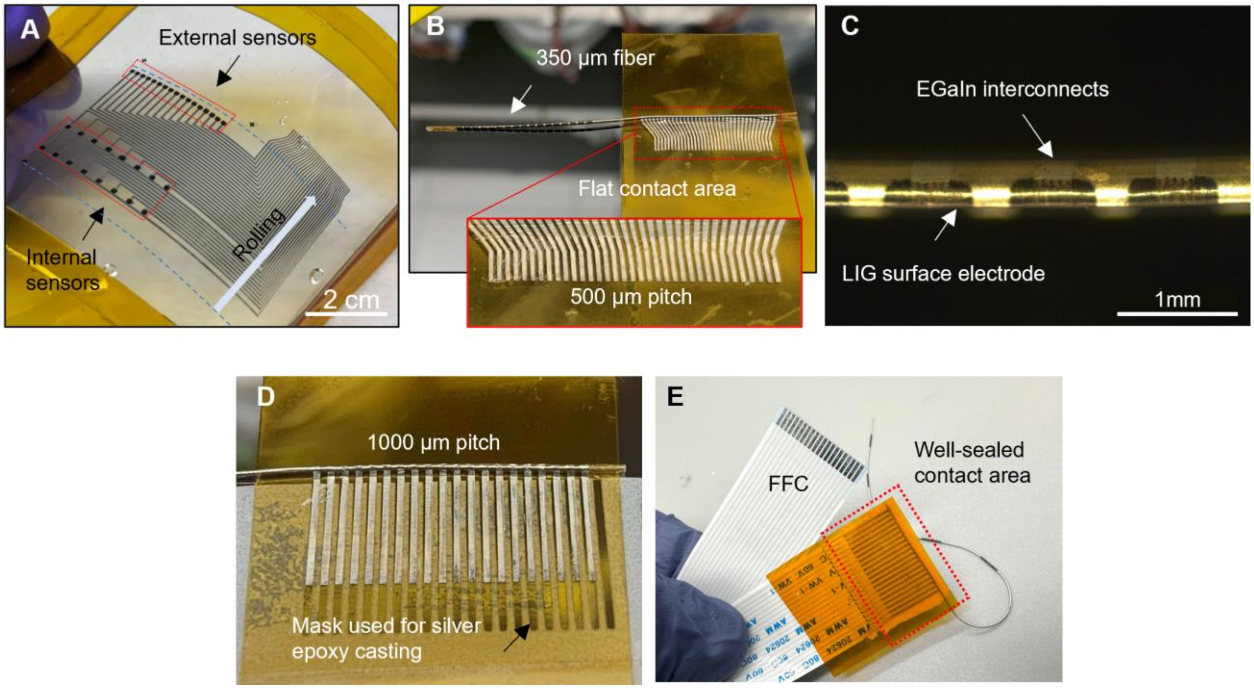


**Fig. S8. S-NeuroString connection to an external flat flexible cable (FFC).** A photograph of a 2D microfabricated film before (**A**) and after (**B**) transformation. The 40 channels (500 µm pitch) are kept unrolled for further connection. The two dashed lines indicate the location of the starting and ending positions of rolling. (**C**) An optical microscope image of the fiber presented in (B). (**D**) A photograph of the mask used to extend the 20 EGaIn channels (1000 µm pitch) with silver paste for further connection with an FFC. (**E**) A photograph showing the bioelectronic fiber connected to an FFC. The connection area is well sealed with hard epoxy (Torr Seal®), which minimizes noise that might be caused by movement or deformation.

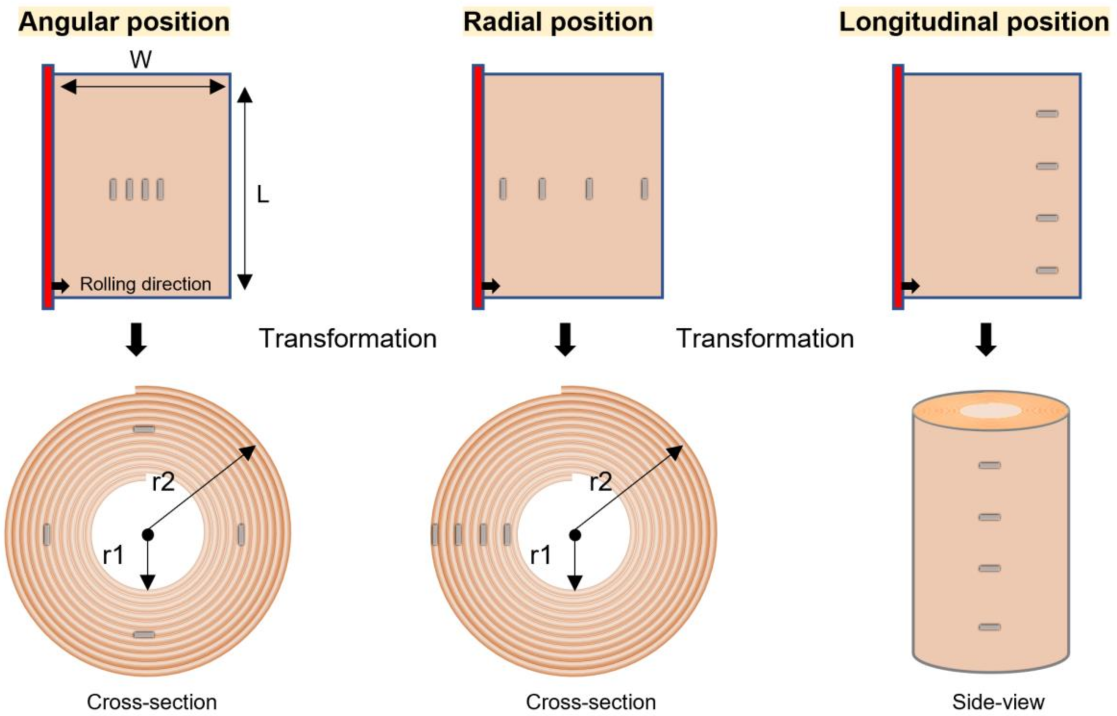


**Fig. S9.** Schematic illustration showing the control over device (a) angular, (b) radial, and (c) longitudinal position.

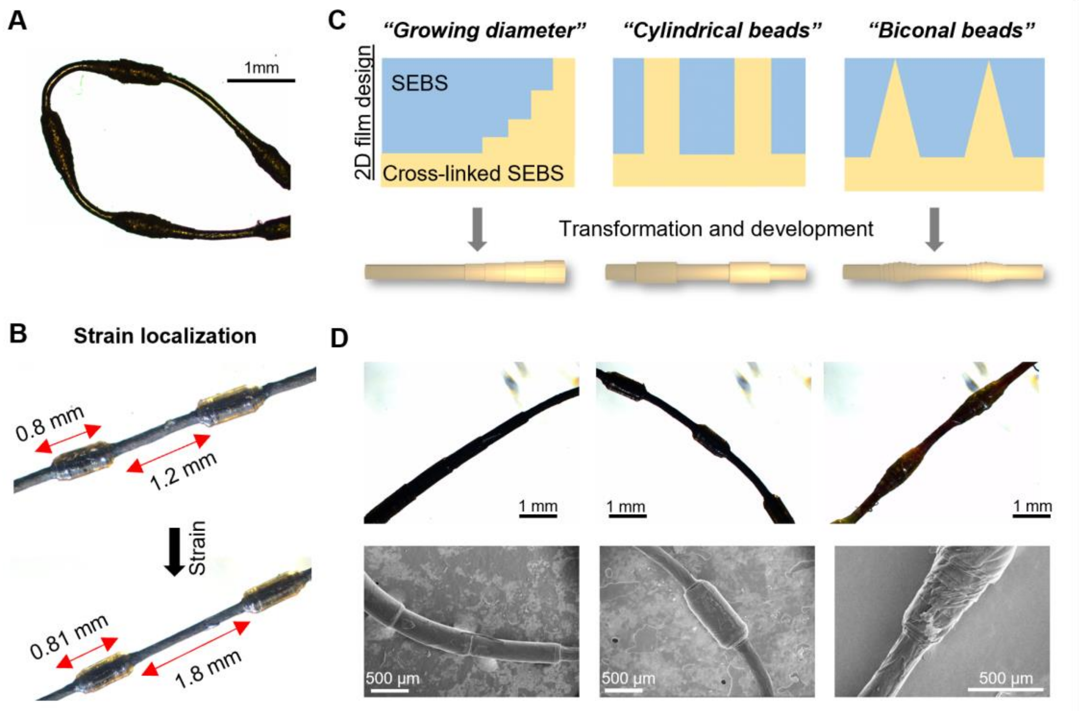


**Fig. S10. Advanced fiber structures and functions.** (**A**) A microscopy image showing a fiber with a periodically changing diameter ('shish-kebab' structure). (**B**) Strain localization could be achieved by using fibers with non-uniform diameter. (**C**) Schematic design of 2D films (before rolling) and corresponding fibers. (**D**) Optical microscope and SEM images of rolled-up and developed fibers. To obtain these fibers, the non-cross-linked SEBS is dissolved after rolling.

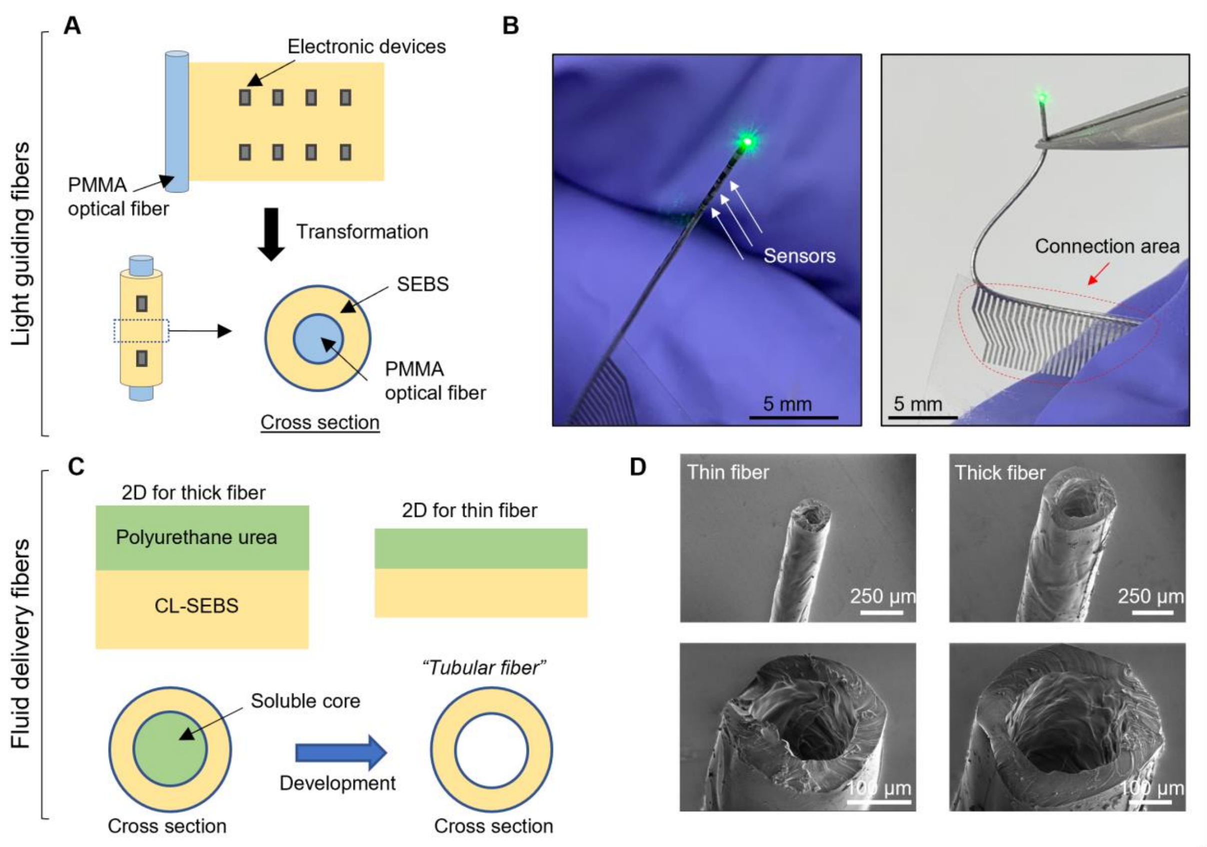


**Fig. S11. Advanced fiber structures and functions.** (**A**) Schematic design of 2D films and fiber rolling using an optic fiber in the core. (**B**) Photographs showing the optoelectronic fibers that can deliver light in addition to its basic electrical capabilities (e.g., sensing). (**C**) Schematic design of 2D film used to prepare hollow fibers. After rolling, the polyurethane urea in the core can be dissolved in acetone to obtain a hollow tube. (**D**) SEM images of hollow fibers.

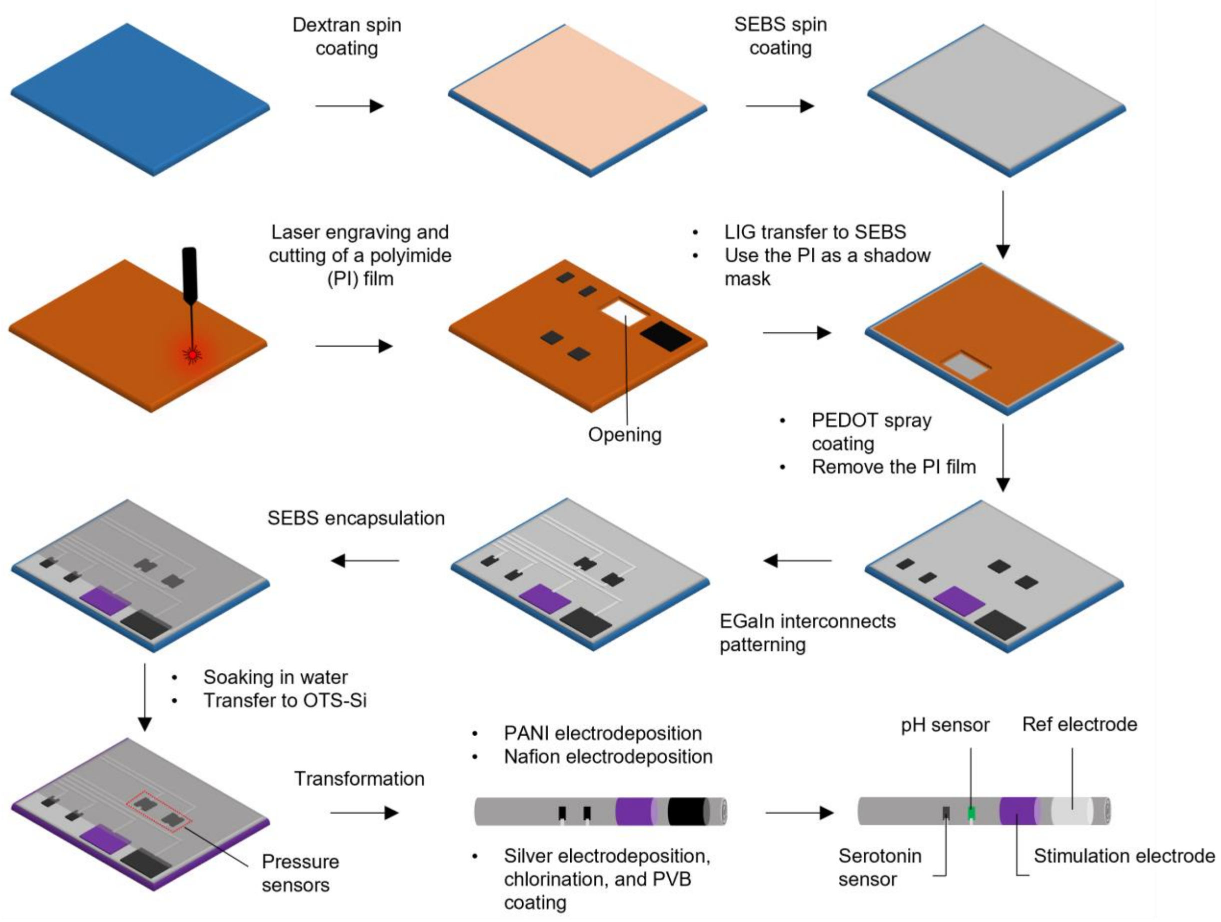


**Fig. S12. Schematic illustration of the microfabrication process used for creating GI S-NeuroString.** LIG is prepared on a polyimide (PI) film using laser writing. Openings are cut on the same PI film by laser. Separately, dextran and SEBS are spin-coated on a Si wafer. LIG is transferred from Kapton to SEBS by sticking the polyimide substrate containing LIG patterns on SEBS and applying pressure. PEDOT-PSS ink is sprayed on the film through the PI shadow mask. This is followed by thermal and UV cross linking. A photoresist is then used to pattern the EGaIn interconnects. The whole device is encapsulated with another SEBS while keeping the electrochemical sensing, stimulation, and reference electrodes exposed. Then, the film is released from the Si wafer in water bath and transferred to an OTS-treated Si wafer. Then, the 2D film is rolled into 1D fiber.

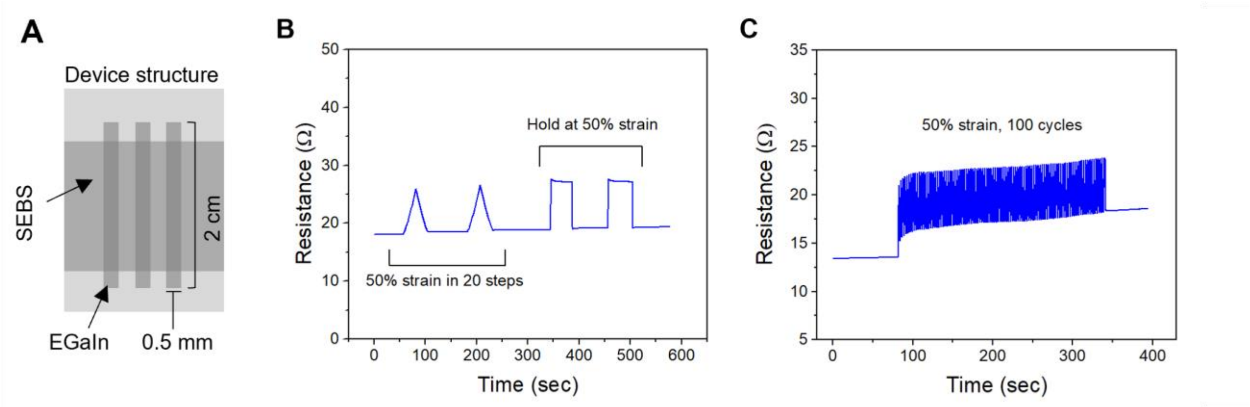


**Fig. S13. Stretchability of SEBS-encapsulated EGaIn interconnects.** (**A**) Schema of the device structure tested in this experiment**.** (**B**) Resistance measurement of prepared EGaIn interconnect under two cycles of stretching to 50% strain (in 20 gradual steps) followed by two cycles of strain hold (at 50 % for 1 min) and release (back to 0 %). (**C**) Resistance measurement of EGaIn interconnects under 100 cycles of 50 % strain.

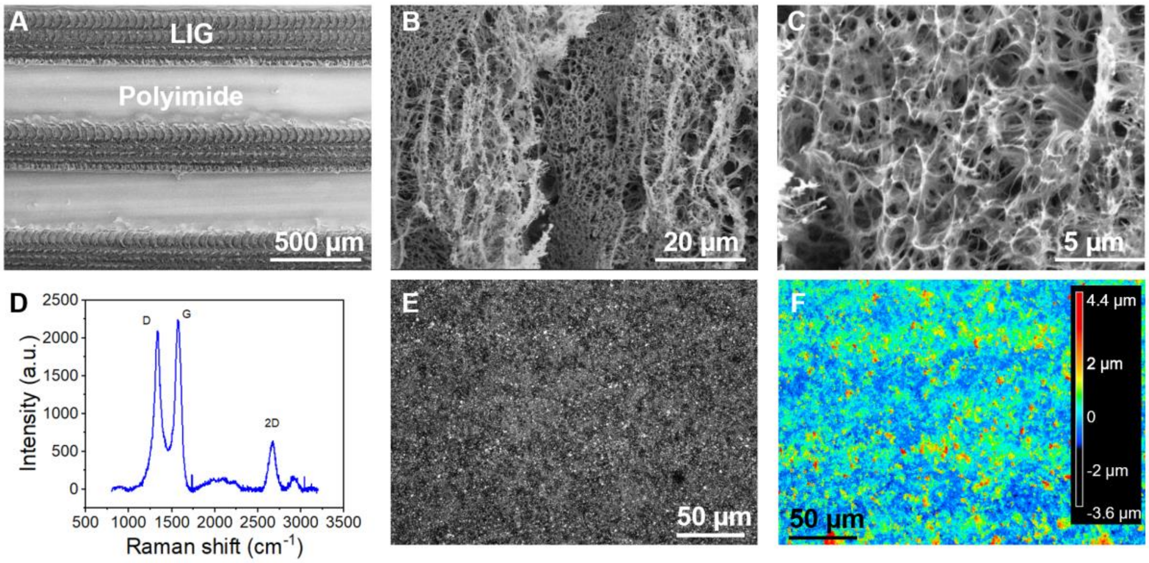


**Fig. S14. LIG used for the development of different sensors.** (**A**)-(**C**) SEM images of LIG fabricated on a polyimide film. (**D**) Normalized confocal Raman spectra (633-nm laser excitation) of our prepared LIG. (**E**) Optical surface image of transferred (from Kapton to SEBS) LIG. (**F**) Height color map showing the surface roughness of the transferred LIG.

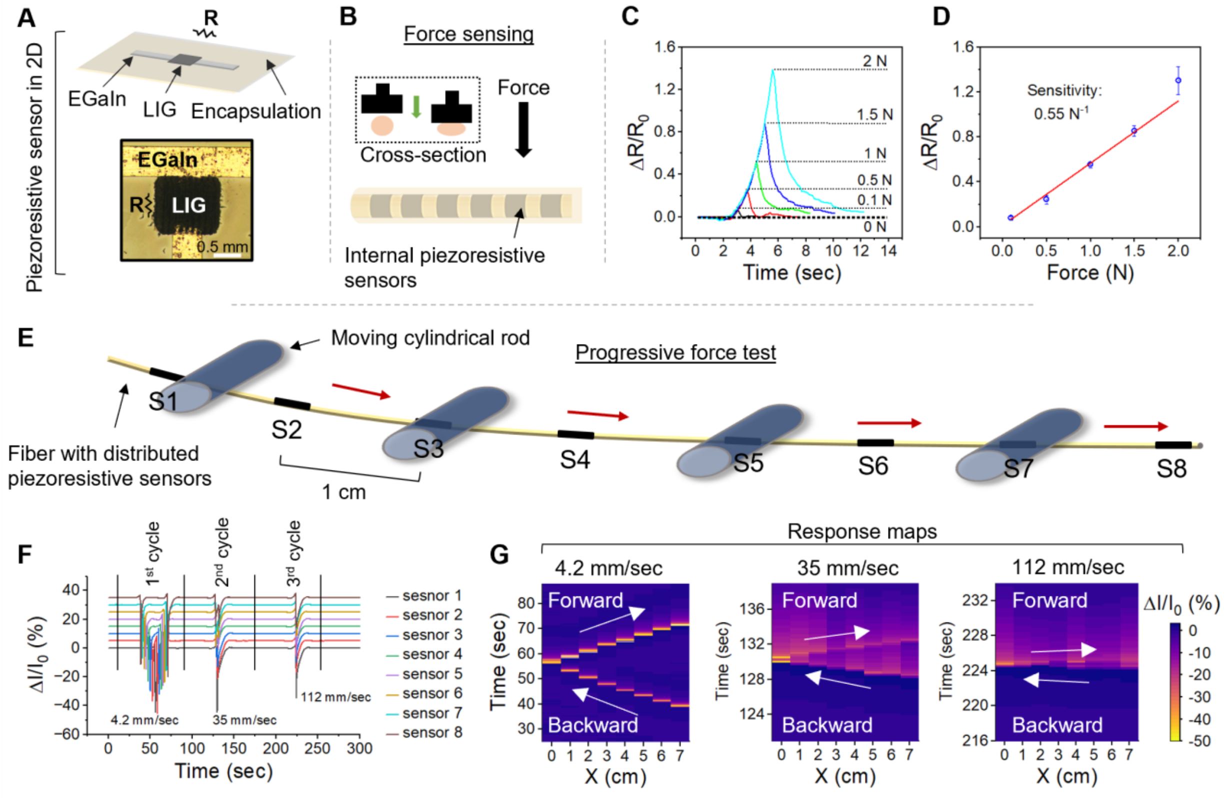


**Fig. S15. Characterization of the pressure and strain sensing.** (**A**) A schematic and microscopic image showing the structure of the LIG-based piezoresistor used for force sensing. (**B**) A schematic picture showing the force testing experiment where force was applied on the fiber (350 µm in diameter) using a force gauge. (**C**) Pressure sensor responses to different applied forces. (**D**) A summary of the sensor's responses to different applied forces showing a good linear sensing behavior (n = 3). (**E**) Schematic illustration of the rolling test used to analyze progressive force sensing. (**F**) The response of 8 sensors to progressive force (starting from sensors #1, moving to sensors #8, and back to sensors #1) with different speeds. All traces are normalized and shifted for clarity (see methods). (**G**) Heat maps showing the response of the sensors to the progressive force described in (F). The sensors could still detect the forward and backward movements even at high progression speed (112 mm/sec).

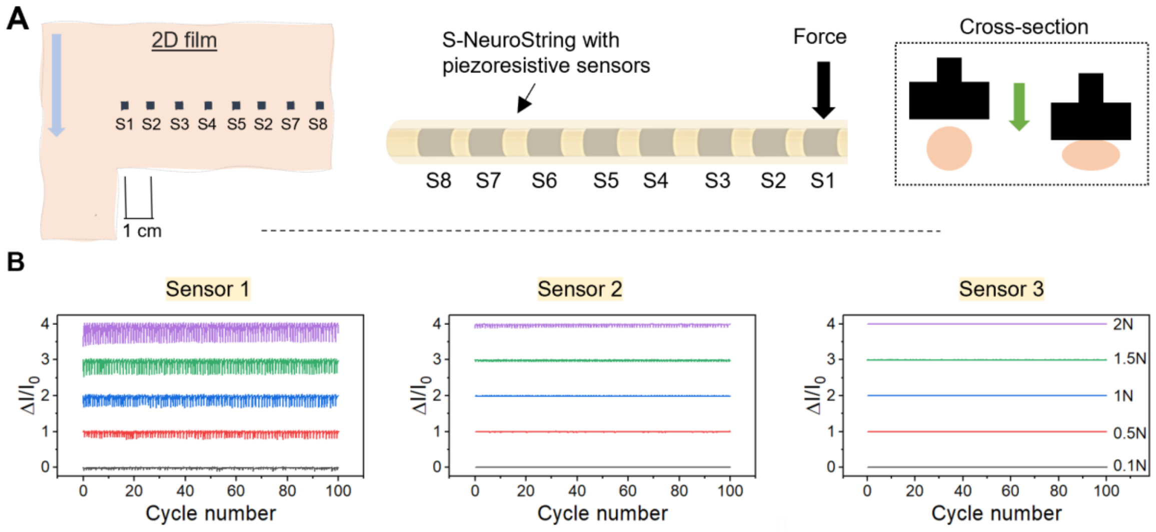


**Fig. S16. S-NeuroString for force mapping.** (**A**) Schematic illustration showing the 2D film containing 8 pressure sensors (S1-S8), corresponding fiber (350 µm), and the test used for force mapping and cycling. (**B**) Force cycling and mapping results: 100 cycles of force-no force were applied on sensors #1. The responses of sensors #1, 2, & 3 were recorded showing good repeatability, spatial selectivity, as well as the insensitivity of EGaIn interconnects to applied forces. All traces are normalized and shifted for clarity (see methods).

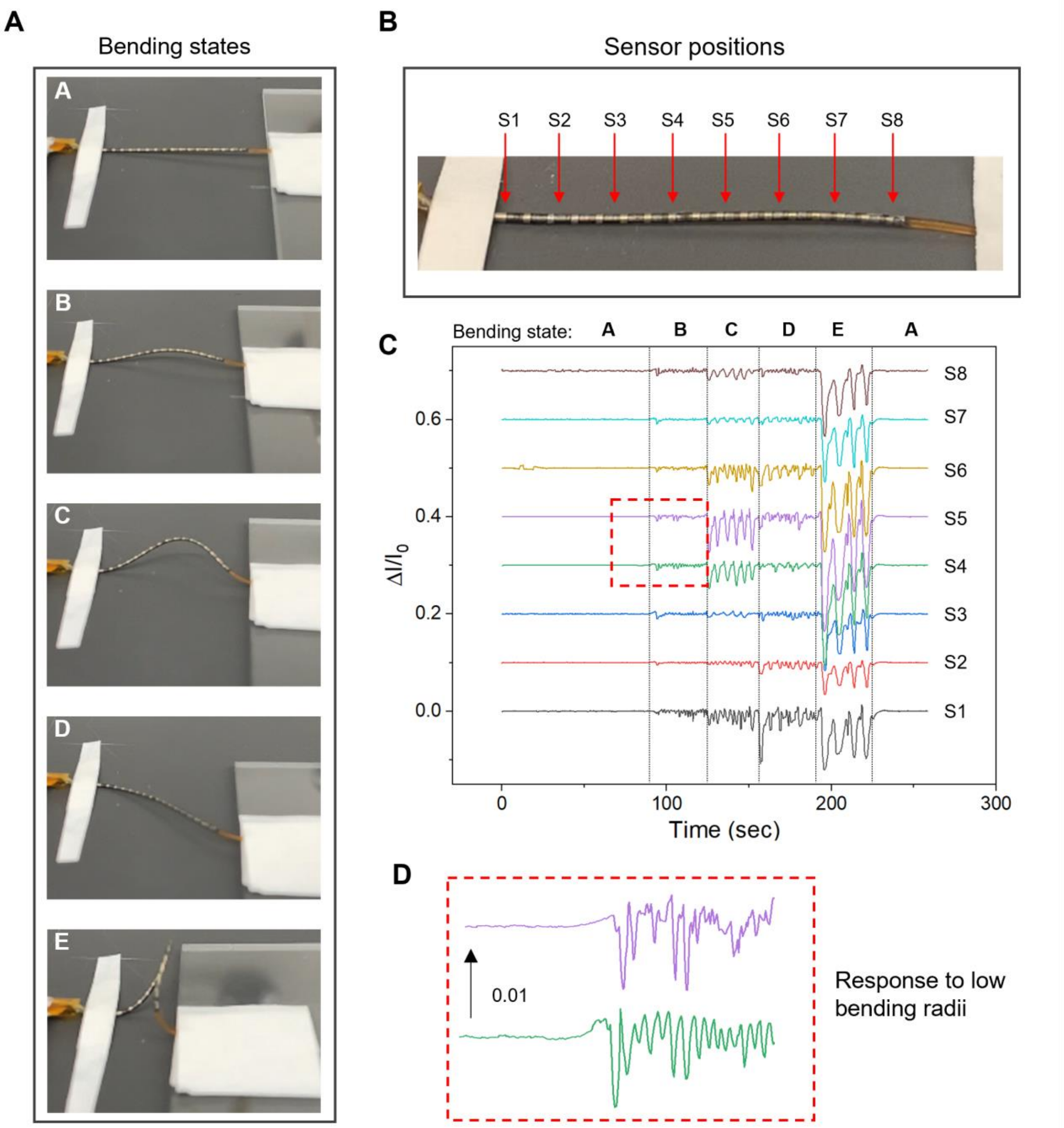


**Fig. S17. Fiber response to different bending states.** (**A**) Photographs showing the different tested bending states. The fiber was manually and continuously transitioned between the original state (A) and the other states (B-E). (**B**) The 8-sensor fiber (350 µm in diameter, 2 cm sensing area) used in this test. (**C**) The response of the 8 sensors to cycles of transitioning from state A to the different other states (B-E). Highest responses are obtained from the middle area (4-5) due to its high curvature compared to the other parts. (**D**) Zoom-in on the tiny responses generated by the transitioning between "A" and "B". All traces are normalized and shifted for clarity (see methods).

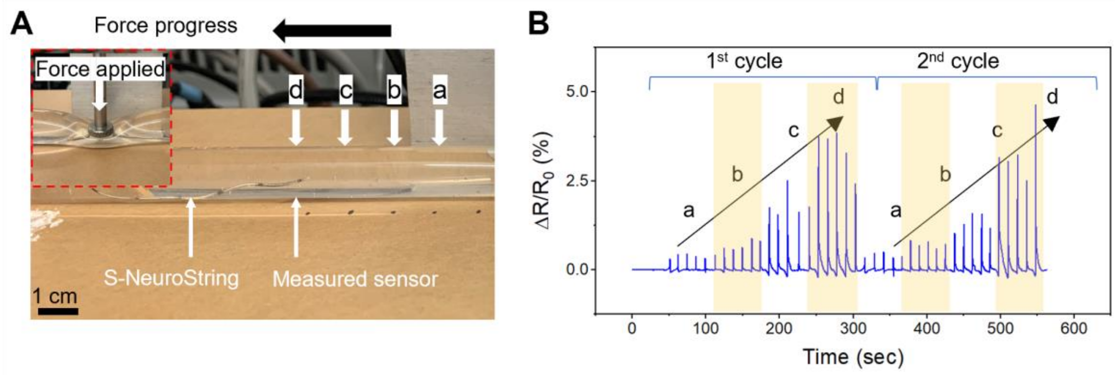


**Fig. S18. In-tube pressure sensing demonstration.** (**A**) A photograph of the system used for testing progressive forces showing the water-filled Tygon tube (Tygon® E-3603; wall thickness: 0.062 in; external diameter: 0.5 in, modulus: 12.1 MPa), S-NeuroString (350 µm in diameter), and force applicator. The tube used to mimic the structure of the intestine. External force (16 N) was applied on the wall of the tube to cause a deformation, similar to the contraction events that may take place in the intestine. Force was applied at different points ("a"-"d") while measuring the response of the sensor located exactly at point "d". (**B**) Sensor's response to forces applied at the different points ("a"-"d"), showing potential for analyzing progressive motility patterns over extended GI segments.

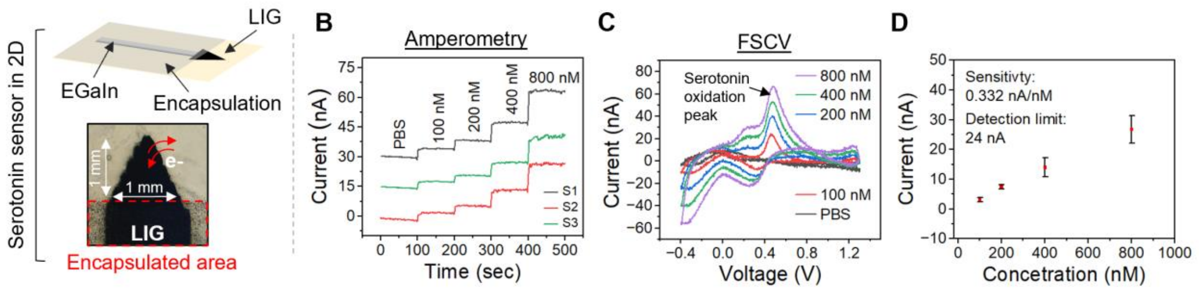


**Fig. S19. S-NeuroString for electrochemical sensing of serotonin.** (**A**) A schematic illustration and microscopy image showing the structure of the electrochemical electrode used for serotonin sensing. (**B**) Amperometric detection of serotonin at 0.5V from 3 different sensors. (**C**) Cyclic voltammetry used for the detection of serotonin, verifying the existence of serotonin peaks. (**D**) A calibration curve showing the response of the sensors to different concentrations of serotonin showing a good linear behavior (n = 4). The slope which represents the sensitivity was calculated from the lowest three concentrations. The limit of detection was calculated based on a signal to noise ratio of 3.

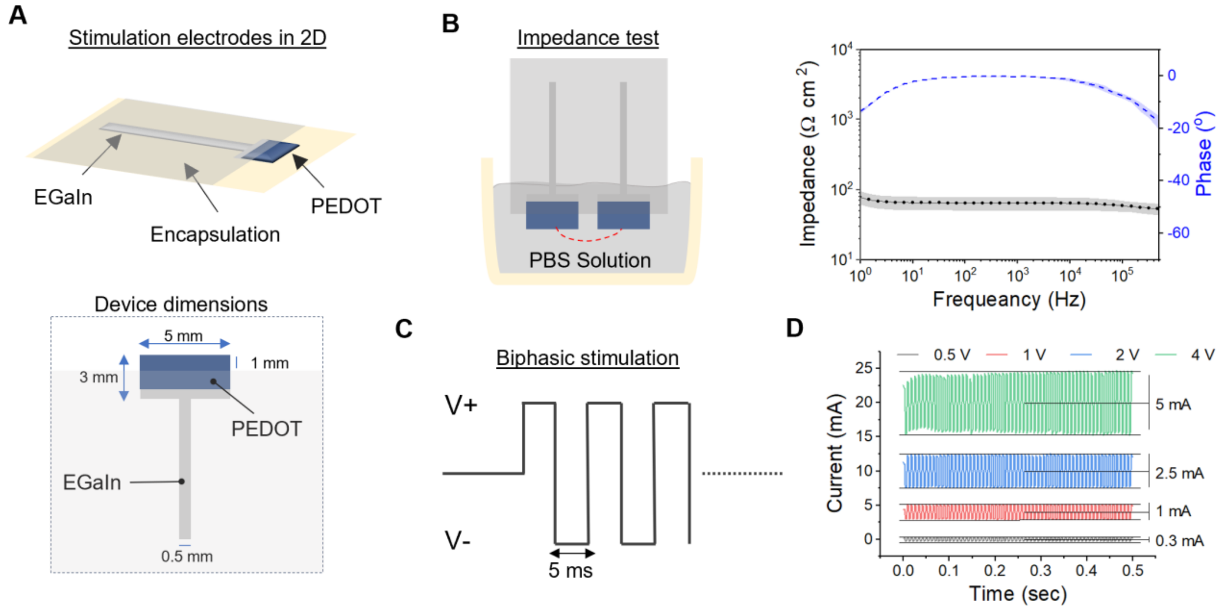


**Fig. S20. Structure and characterization of the PEDOT-PSS based stimulation electrodes**. (**A**) Schematic illustrations showing the structure and exact dimensions of the PEDOT-PSS based stimulation electrode in 2D. After rolling, the electrodes ended up as annular rings on the fiber. (**B**) Impedance measurement of the stimulation electrodes in PBS. The shaded area in the graph represents the standard deviation (n=3). (**C**) The parameters of the biphasic stimulation used to trigger motility both in mice and pigs. (**D**) stimulation currents achieved under different voltages in PBS solution.

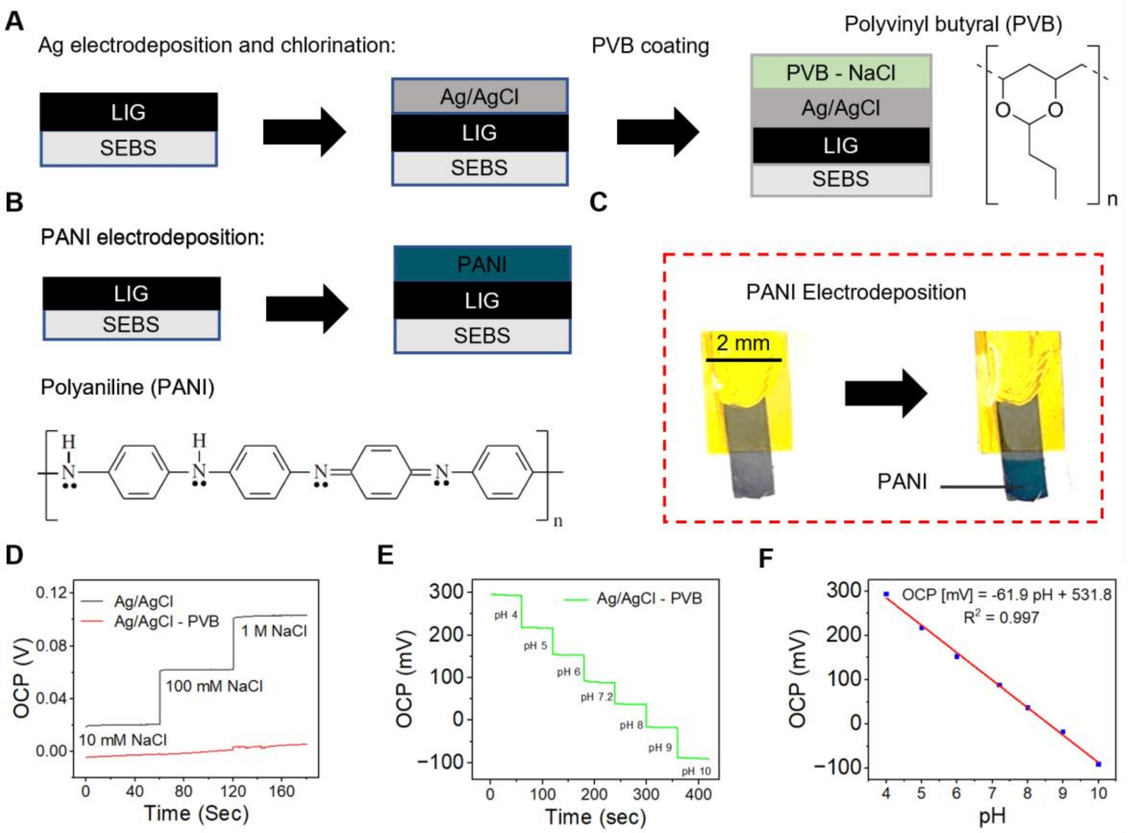


**Fig. S21. Fabrication and characterization of pH sensors**. (**A**) Scheme showing the structure of the reference electrode which is made of PVB-encapsulated AgCl-LIG electrode. (**B**) Scheme showing the structure of the pH sensing electrode which is made of PANI-modified LIG electrode. (**C**) A photograph showing the change in color of an LIG electrode following the electrodeposition of PANI. (**D**) Stability of the PVB-encapsulated vs exposed AgCl-LIG electrode in solutions with different ion concentrations. (**E**)-(**F**) Sensor response to different pH values.

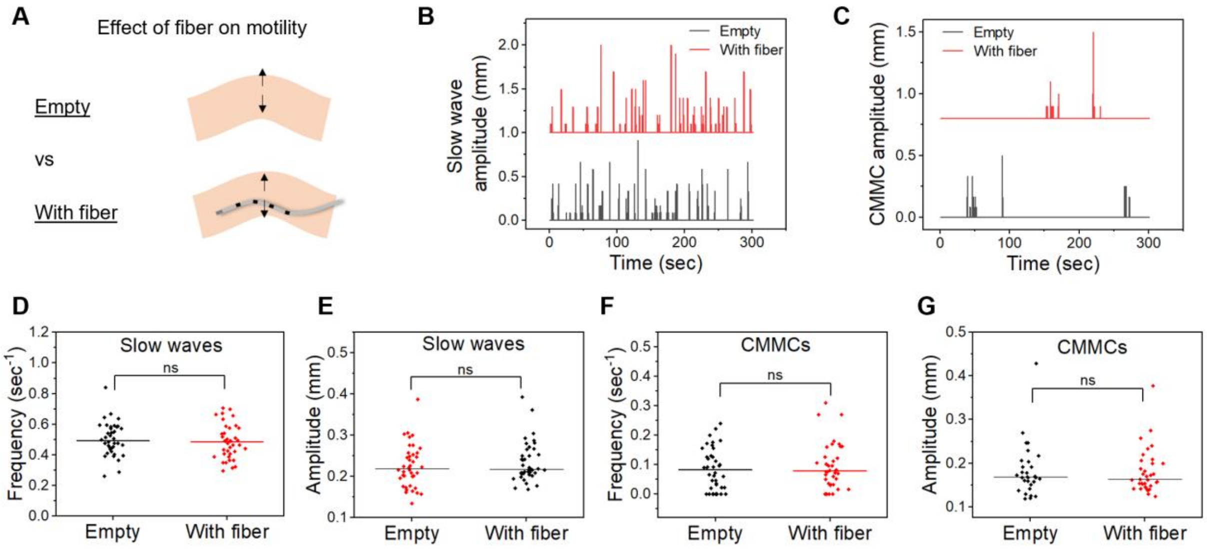


**Fig. S22.** ***Ex-vivo*** **study of the effect of fibers on colonic motility.** (**A**) A schematic picture showing the ex-vivo experimental set-up used to study the effect of fibers on colonic motility. Motility signals (calculated from video analysis) were compared before and after the insertion of S-NeuroString inside. (**B**) Detected slow motility wave events in empty colons and colons with a bioelectronic fiber inserted inside. (**C**) Detected CMMCs events in empty colons and colons with a bioelectronic fiber inserted inside. (**D**)-(**G**) Frequency and amplitudes of slow wave and CMMC events in empty colons and colons with a bioelectronic fiber inserted inside. (D)-(G), Each group contains 40 data points obtained from colonic diameter monitoring at 8 different locations in 5 mice.

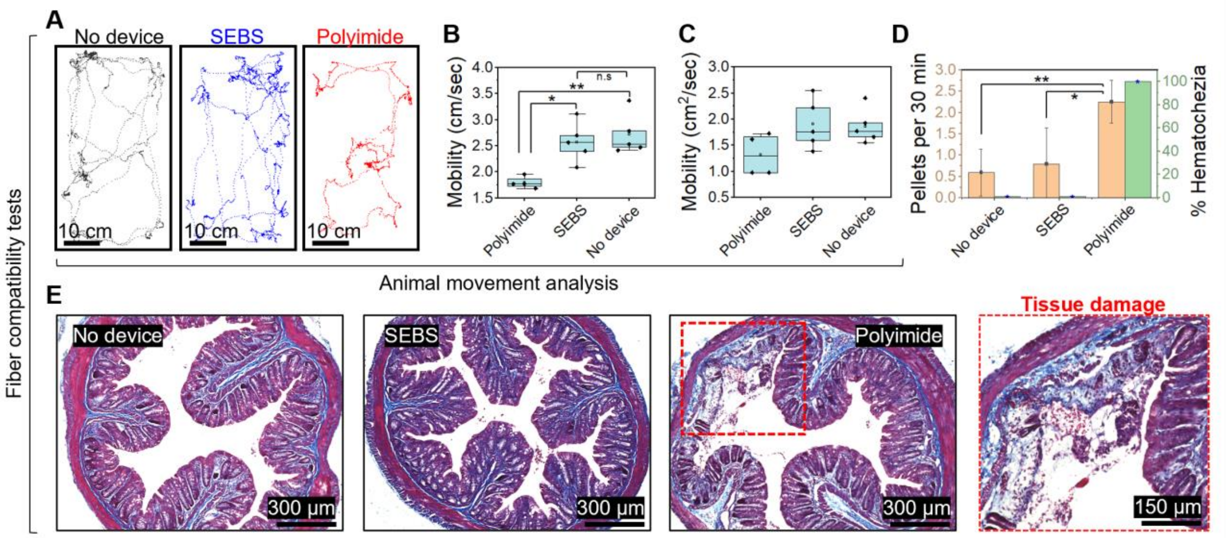


**Fig. 23. Fiber compatibility in awake mouse.** (**A**)-(**C**)**,** Movement analysis of mice with SEBS (H1062) fiber (300 µm in diameter and 5 MPa Young's modulus) or a flexible polyimide (300 µm wide, 127 µm thick) inserted in the colon during the first 30 min after insertion (n = 5). (**D**)**,** Pellet output and hematochezia rate for mice implanted with an SEBS fiber and a flexible polyimide during the first 30 min after insertion (n = 5). P value: ns P > 0.05; * P ≤ 0.05; ** P ≤ 0.01; *** P ≤ 0.001; unpaired, two-tailed Student's t-test. (**E**)**,** Representative Trichrome staining images of colon tissues collected from freely moving animals after 30 min of insertion from the three groups described previously (sham, SEBS, and polyimide). The right image shows a zoom-in on the damage caused by the existence of polyimide in the colon.

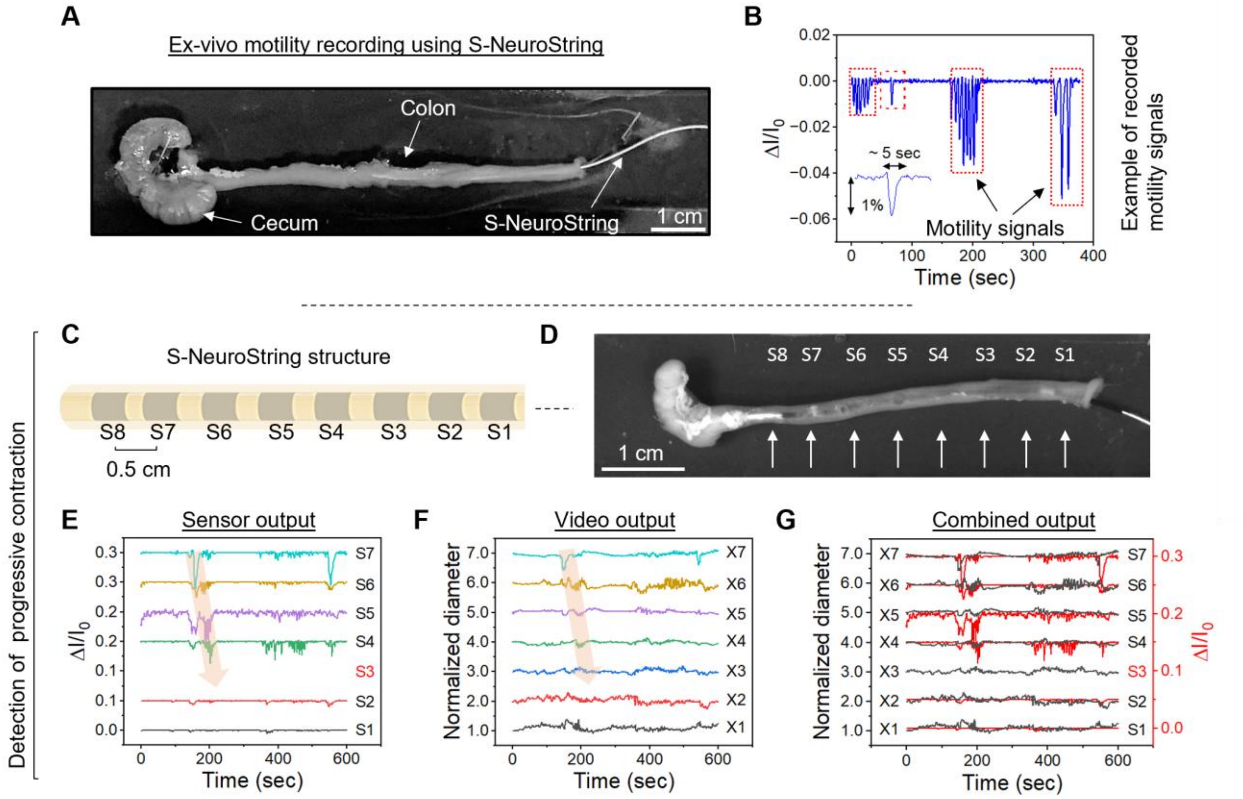


**Fig. S24.** ***Ex-vivo*** **colonic motility sensing.** (**A**) A photograph showing the system used for ex-vivo monitoring of colonic motility. (**B**) Motility signals recorded by S-NeuroString (350 μm in diameter, 5 MPa Young's modulus) inside the colon. (**C**) Structure of the S-NeuroString used in the progressive motility detection test. (**D**) A Photograph showing the colon from Movie S2 and the location of the different sensors used for motility detection. The distance between each two sensors is 0.5 cm. (**E**) Motility signals recorded by sensors in the colon. Progressive pressure signals were observed both in the sensors' output (marked with an arrow) and Movie S2. (**F**) Measured diameter recorded by a camera at different locations on the colon (X1 – X7). S1 is located at X1, S2 is located at X2, etc. These results show the same progressive wave that appears in (**E**), which starts at t = 150 sec. (**G**) Comparison between the sensor output and video analysis output, showing very high similarity between the signals, which further proves the precise motility sensing provided by our fibers. All traces are normalized and shifted for clarity (see methods).

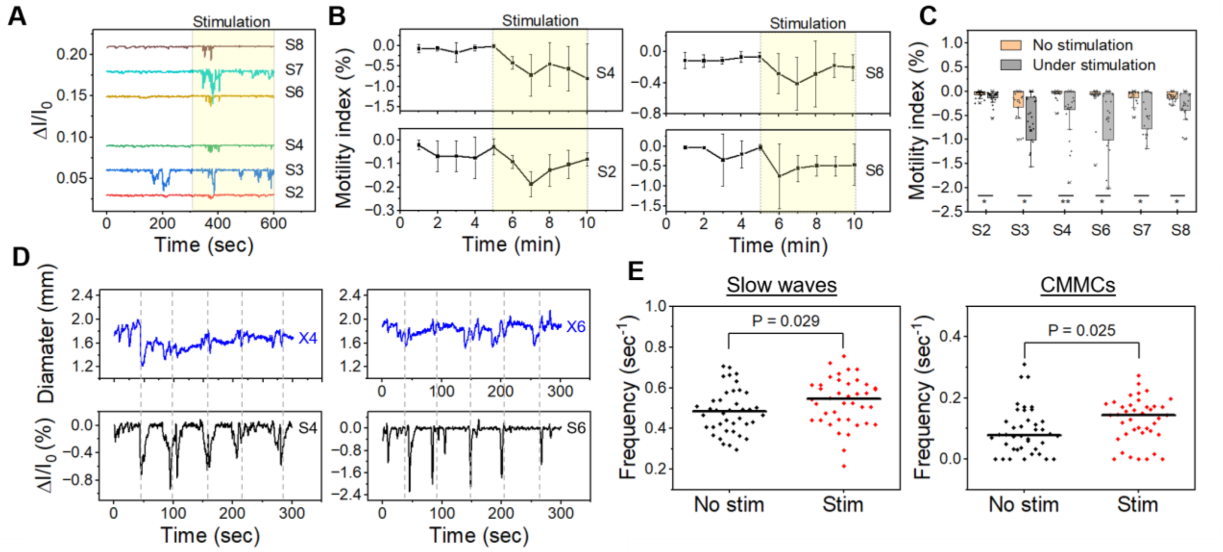

**Fig. S25.** ***Ex-vivo*** **colonic motility sensing and stimulation.** (**A**) Output of piezoresistive sensors before and during electrical stimulation of the tissue (1.5 mA, 100 Hz). All traces are normalized and shifted for clarity (see methods). (**B**) Time course of the motility index obtained by different piezoresistive sensor (S2, S4, S6, S8) before and during tissue stimulation. Under stimulation, 5 stimulation steps were performed. Each stimulation step (1.5 mA, 100 Hz) lasted for 5 seconds and followed by recovery time (no stimulation) for 55 seconds. This was repeated on 5 colons (n=5 mice). **(C)** Summary of motility index values before and during stimulation, which is calculated from the output of the different sensors (n = 5 mice). P value: ns $P > 0.05$; *$P \leq 0.05$; **$P \leq 0.01$; ***$P \leq 0.001$; paired, two-tailed Student's t-test. (**D**) Overlay of sensor output and measured diameter as a function of time. Dashed lines mark the beginning of stimulation. S4 and S6 are located at X4 and X6, respectively. (**E**) Frequency of slow wave and CMMC events before and after stimulation. Each group contains 40 data points (5 mice and 8 different locations on the colon).

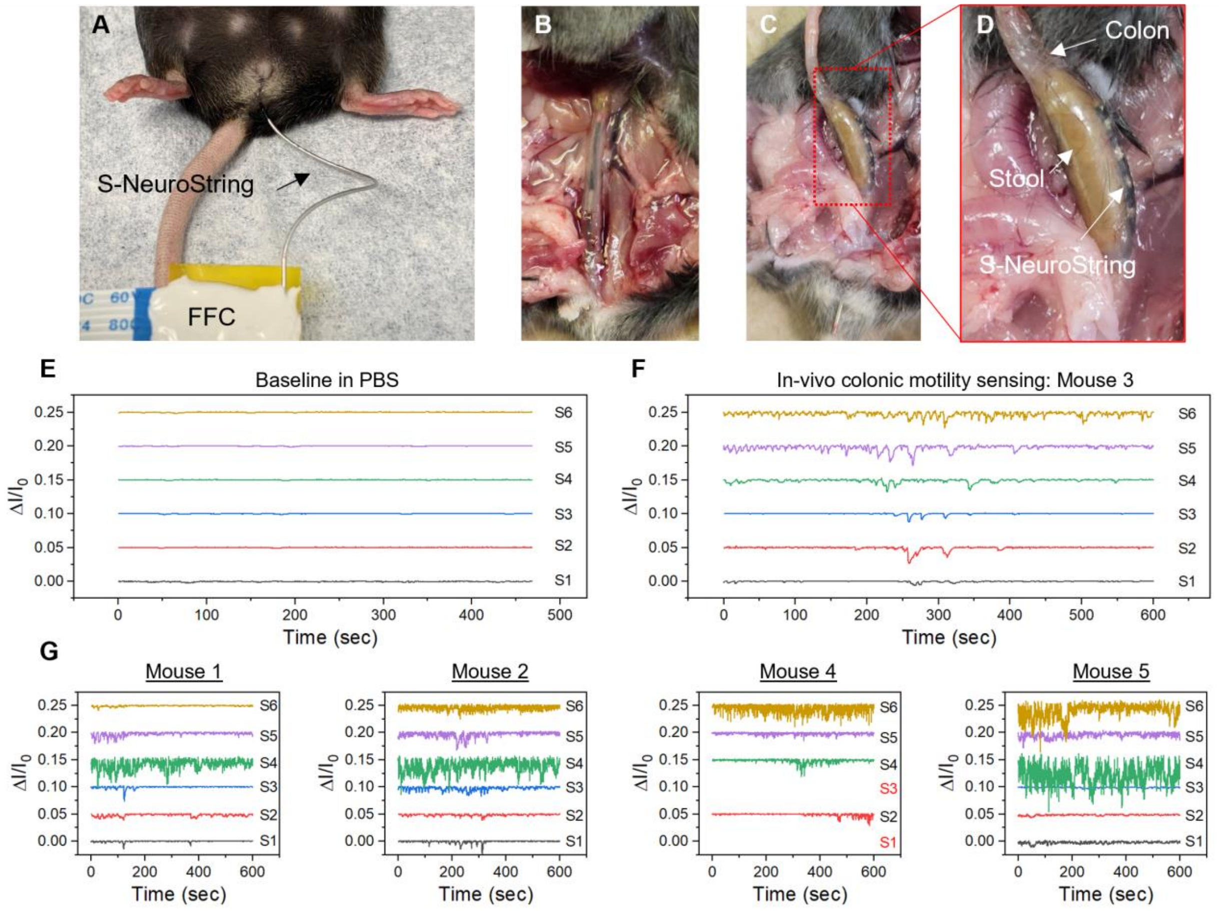


**Fig. S26.** (**A**) A photograph showing the bioelectronic fiber inserted in the colon of an anesthetized mouse. (**B**)-(**D**) Photographs showing an exposed colonic segment with a S-NeuroString inside. (**E**) Measurements obtained from 6 sensors located on S-NeuroString in PBS solution, showing stable sensor performance under aqueous conditions. (**F**) Motility recording obtained from the same 6 sensors in the colon of an anesthetized mouse. (**G**) Motility recording in the colons of 4 different anesthetized mice. All traces are normalized and shifted for clarity (see methods). S1 and S4 in mouse 4 are broken sensors.

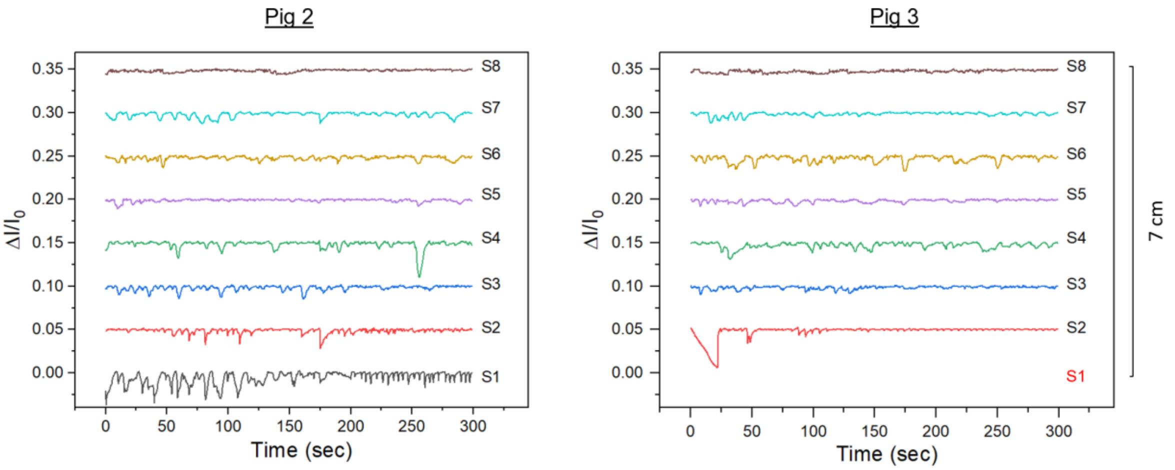


**Fig. S27.** Recording of natural motility in the small intestine of pig using a monofunctional S-NeuroString with 8 pressure sensors. The distance between adjacent sensors is 1 cm. S1 in Pig 3 is broken. All traces are normalized and shifted for clarity (see methods).

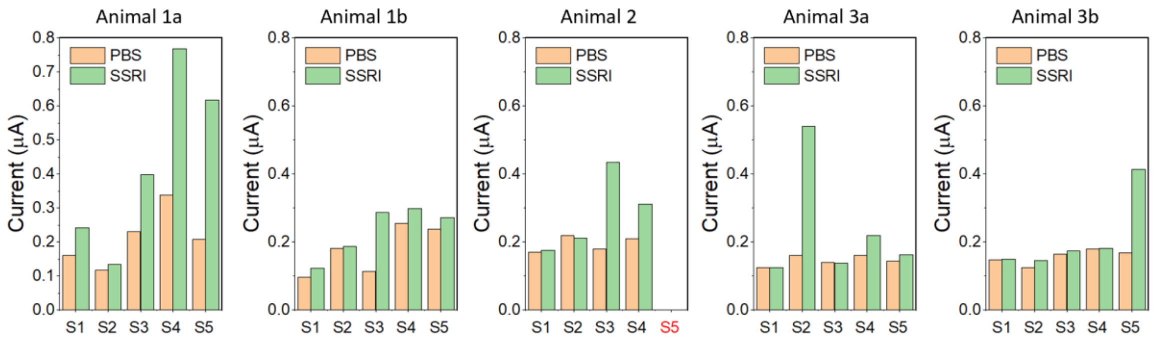


**Fig. S28.** Responses of the different electrochemical sensors after PBS and fluoxetine (SSRI) injections in the small intestine.

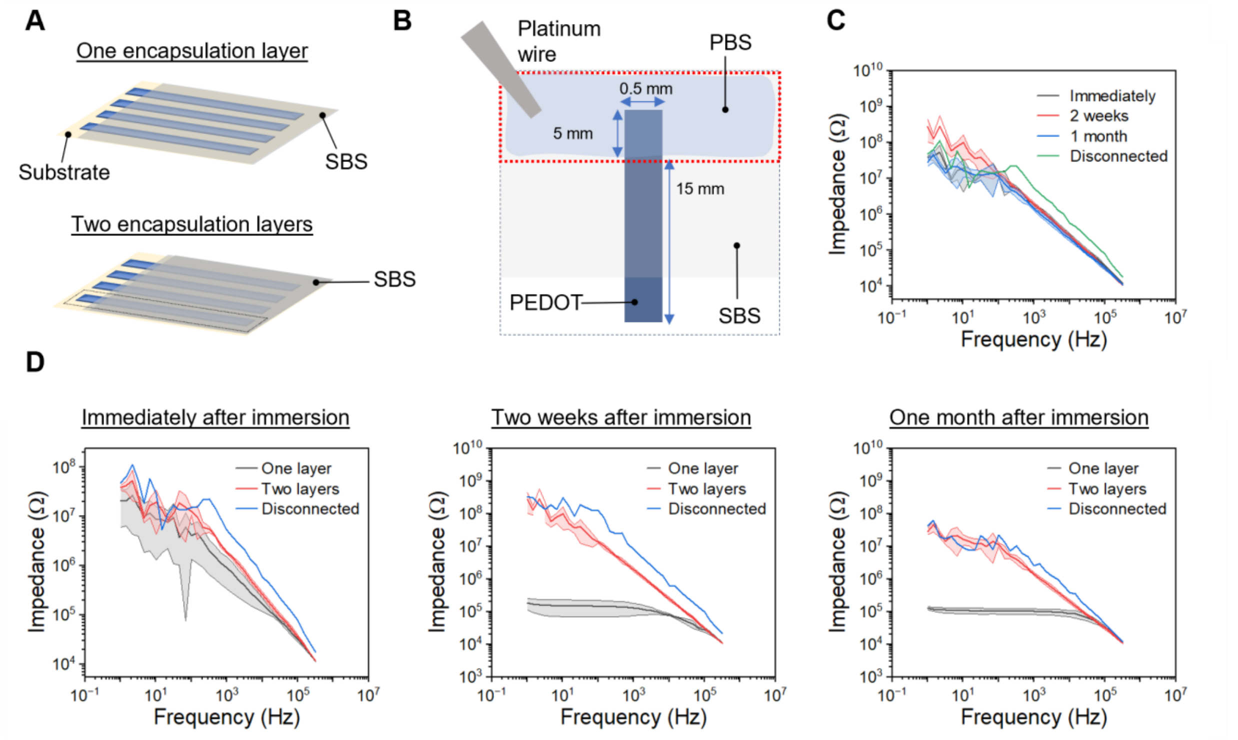


**Fig. S29. SBS encapsulation stability in PBS.** (**A**)-(**B**) Schematic illustrations showing the device structure used for testing the SBS encapsulation stability under PBS. One (1.6 µm) and two (3.2 µm) layers of SBS encapsulation were tested. (**C**) Impedance measurement of double-encapsulated PEDOT electrodes with time. Disconnected represents a control measurement without a device being connected. (**D**) Impedance measurements of single- and double-encapsulated PEDOT electrodes. Shaded areas in c and d represent the standard deviation, n = 4).

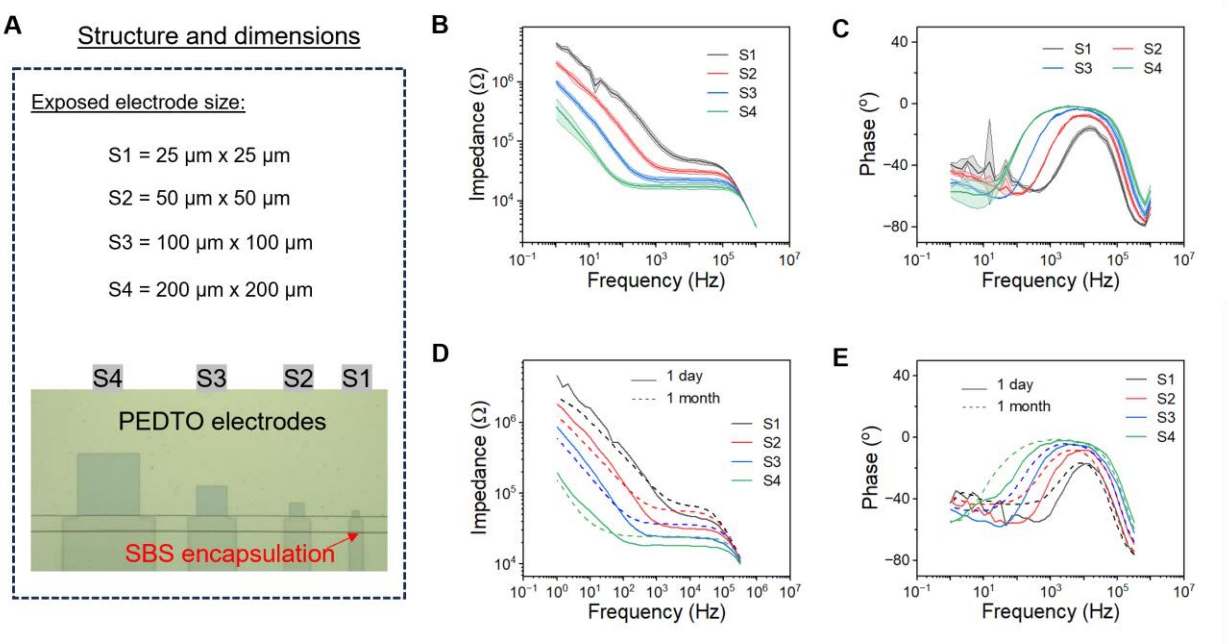


**Fig. S30. PEDOT-based electrodes impedance test**. (**A**) Schematic illustration showing the device structure used for in-vitro impedance tests. Four sizes of PEDOT recording electrodes were prepared: S1, S2, S3, and S4. (**B**)-(**C**) Impedance and phase as a function of size (n=3, shaded area represents the standard deviation)**.** (**D**)-(**E**) Impedance and phase as a function of size after 1 day and 1 month of immersion in PBS solution**.**

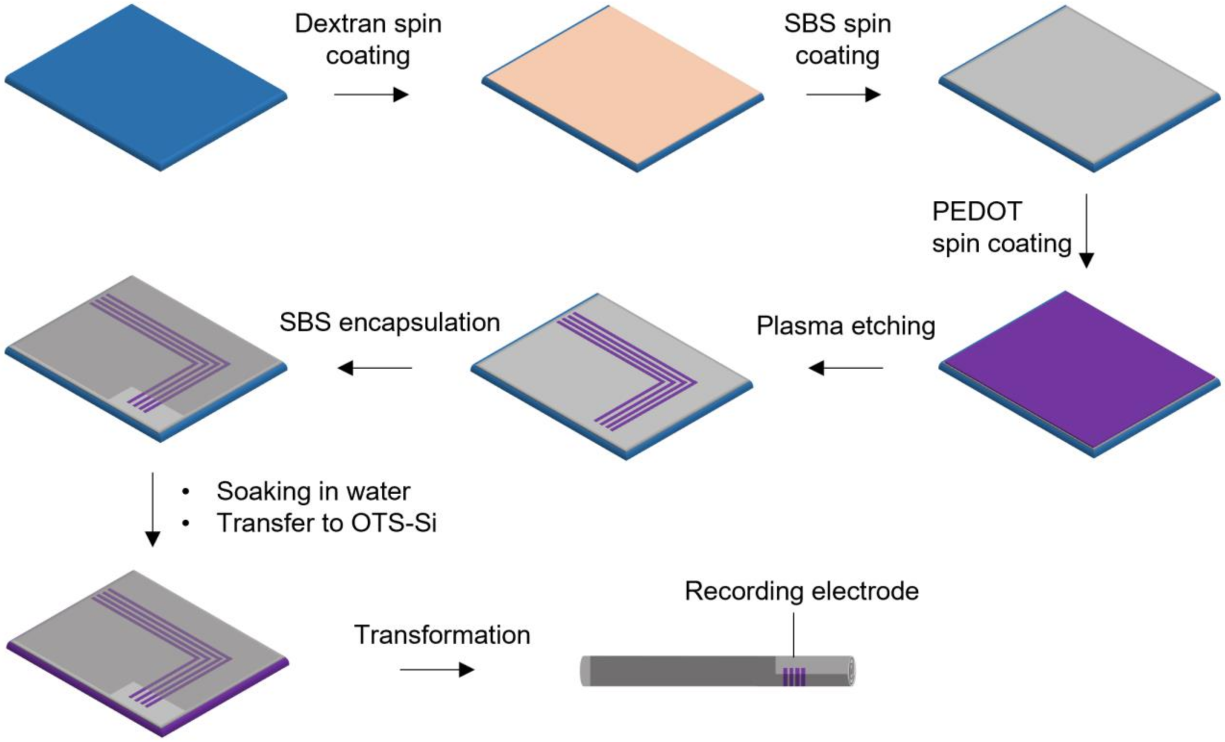


**Fig. S31.** Fabrication process used for creating PEDOT-based recording electrode.

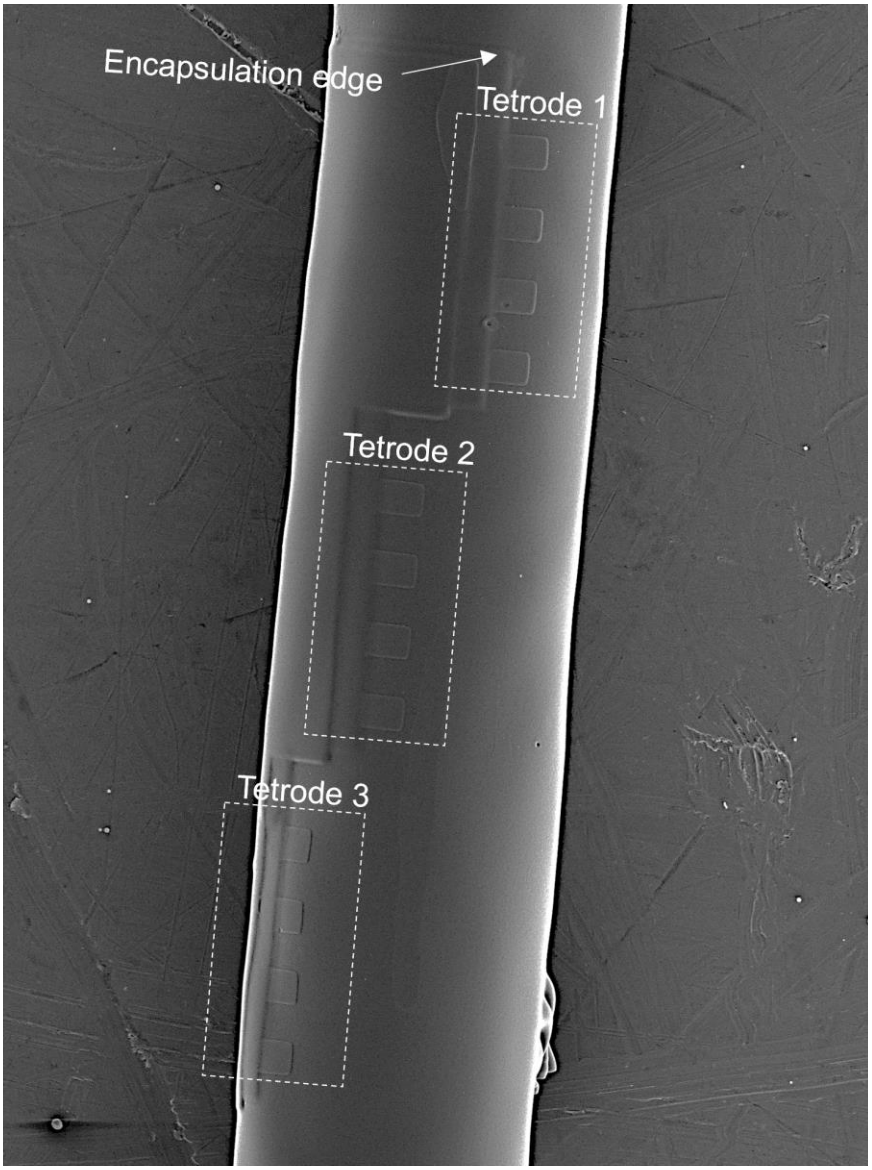


**Supplementary Fig. S32|** SEM image showing the helical distribution of tetrodes at the recording area of S-NeuroString

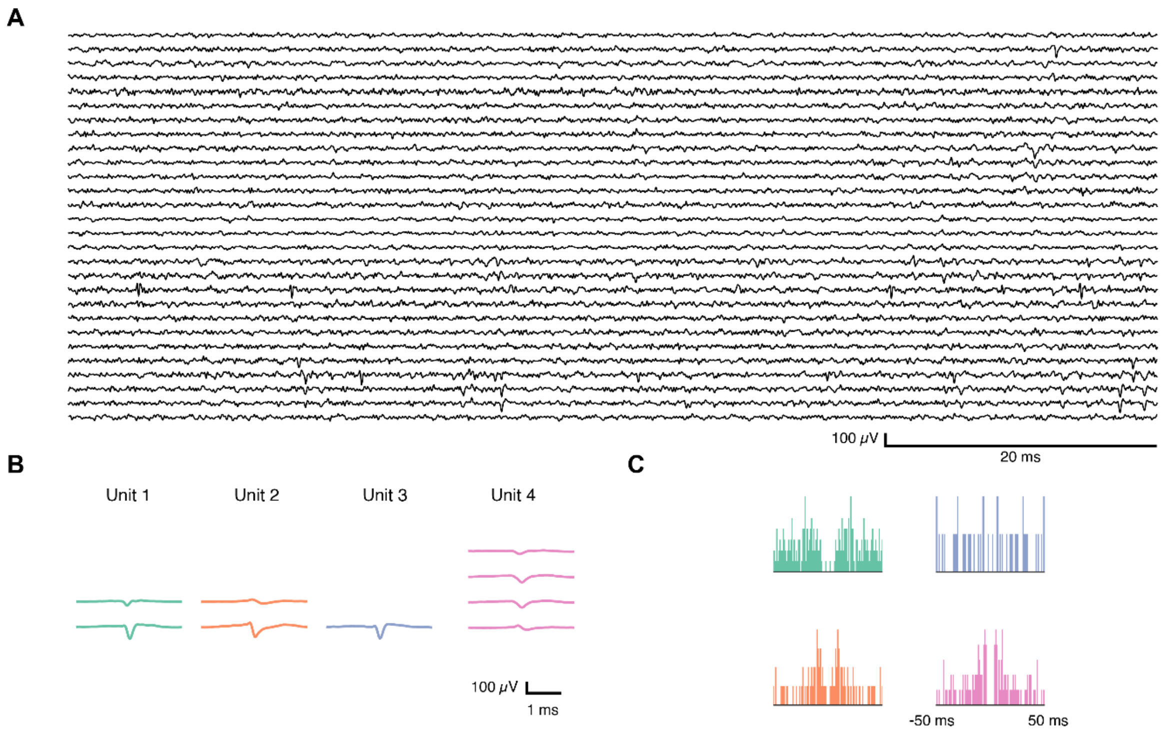


**Fig. S33.** (**A**) S-NeuroStrings can detect spontaneous spiking activity as early as two days post-implantation in the hippocampus as shown with the waterfall plot of the electrophysiological data band-passed at 300 – 7000 Hz. (**B**) The footprint of individual neurons. (**C**) Autocorrelogram of the respective neurons.

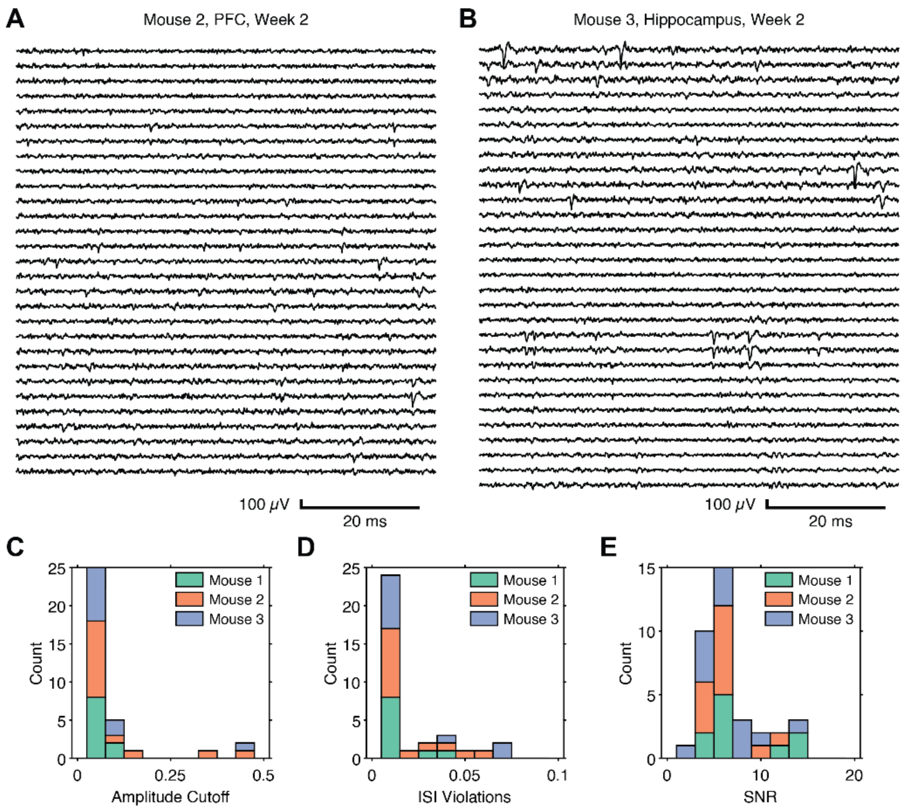


**Fig. S34.** (**A**)-(**B**) Electrophysiological data of two additional S-NeuroStrings implanted in the PFC and hippocampus 2 weeks post implantation respectively, band-passed at 300 – 7000 Hz. (**C**), (**D**), and (**E**) Quality metrics including the amplitude cutoff, ISI violations and SNR indicating that S-NeuroStrings are able to detect high quality single units in multiple regions of the brain. Mouse 1 was implanted in the hippocampus and the data was recorded 1-week post-implantation. Mouse 2 was implanted in the PFC and the data was recorded 2-weeks post-implantation. Mouse 3 was implanted in the hippocampus and the data recorded 2-weeks post-implantation.

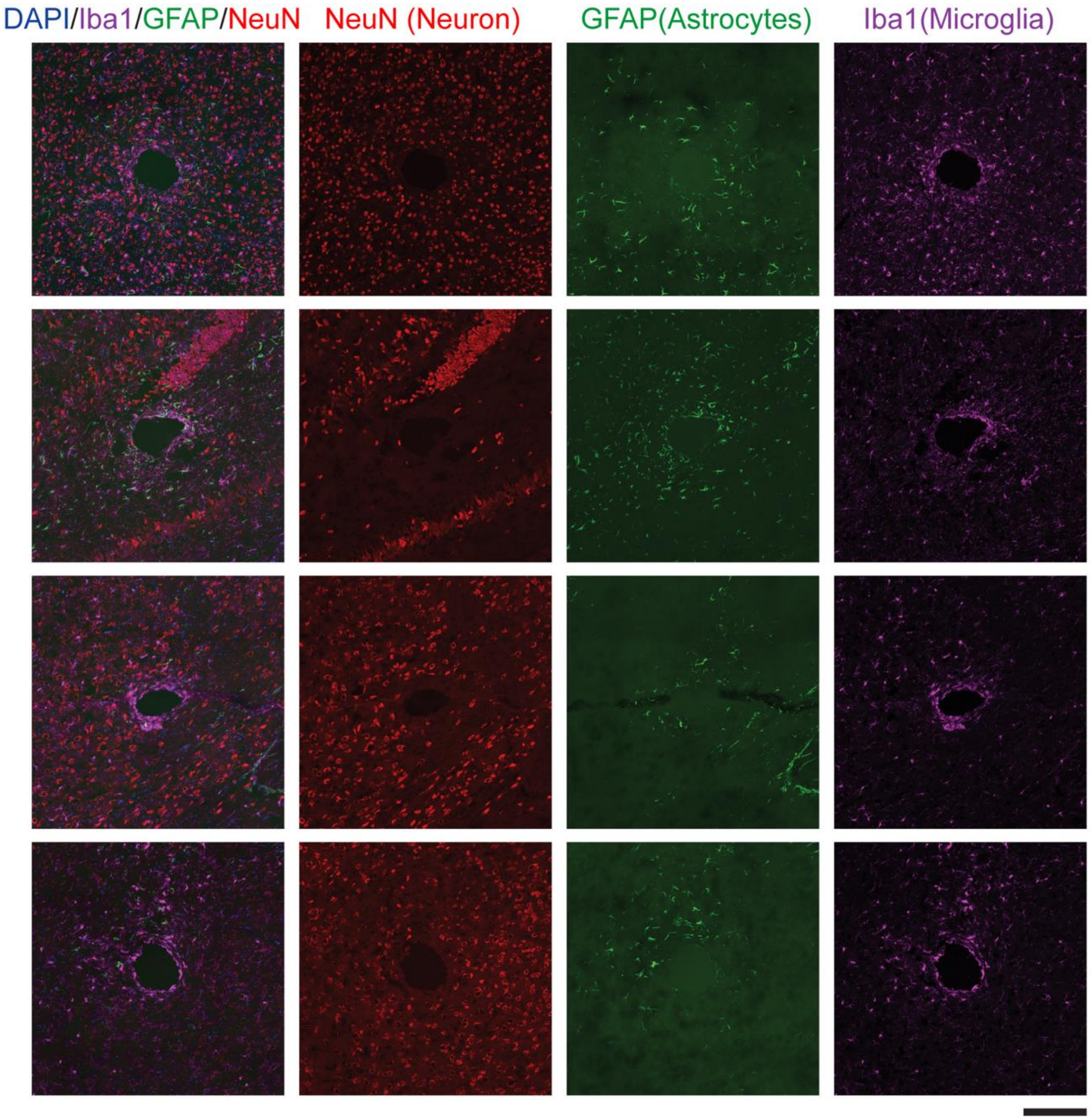


**Fig. 35. Histological investigations of the brain tissue's response to implanted soft fiber electrodes.** Immunofluorescence images depict the tissue reactions after a 1-week implantation of a 150-μm diameter soft fiber electrode. Tissue labeling involved microglia (Iba1, shown in purple), astrocytes (GFAP, green), neurons (NeuN, red), and DAPI staining (blue) (n=4, 3 mice, each row represents one repetition**)**. Scale bar: 200 μm.

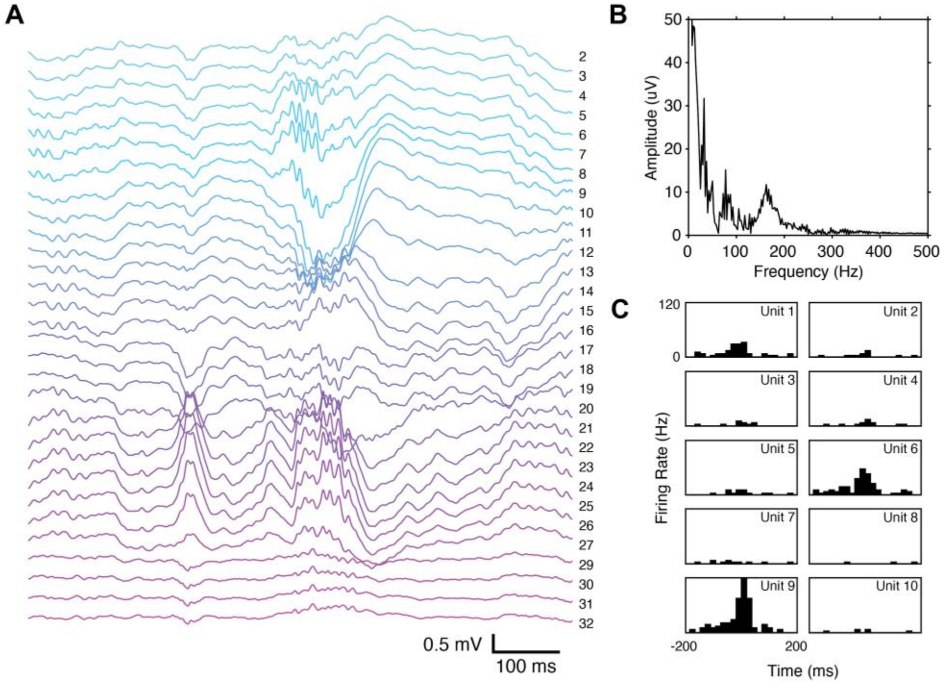


**Fig. S36.** (**A**) Waterfall plot of the raw LFP data recorded one week after a S-NeuroString was implanted in the hippocampus, band-passed Butterworth filtered between 0.1 – 300 Hz. Colors correspond to the depth of the electrodes where blue is more ventral and purple more dorsal. SWR electrophysiological signatures are apparent and consistent with literature (*19*), most notably with the co-occurrence of CA1 and CA3 ripple events as well as an inverse of polarity with depth. (**B**) Discrete Fourier transform of channel 7, with a clear peak at 160 Hz indicative of the SWR. (**C**) Firing patterns of individual neurons around SWRs, averaged across 14 events. Multiple neurons show elevated firing rates around a SWR event.

### Supplementary Sections

#### Supplementary Section S1: Fiber diameter calculation

The diameter of a transformed fiber is determined by the thickness and width of the corresponding 2D film. For the calculations, we assume ideal rolling, i.e., no gaps between the different layers in the rolled structure.

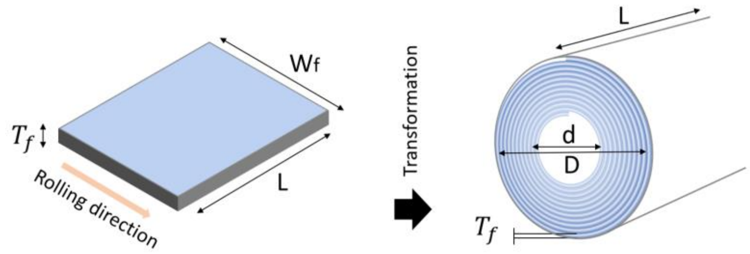


The connection between the dimensions of the unrolled 2D film and the final fiber can be determined by equating the volumes of the 2D film and the transformed fiber:

$$(1) \qquad \frac{\pi L\,(D-d)^2}{4} = T_f W_f L$$

$$(2) \qquad D^2 = \frac{4T_f W_f}{\pi} + d^2$$

$$(3) \qquad D = \sqrt{\frac{4T_f W_f}{\pi} + d^2}$$

### Supplementary Section S2: Density gain following the transformation process

To evaluate the gain in density that can be obtained by using the rolling process, we compare the component density on a rolled fiber and the corresponding rectangular 2D film as shown in the following:

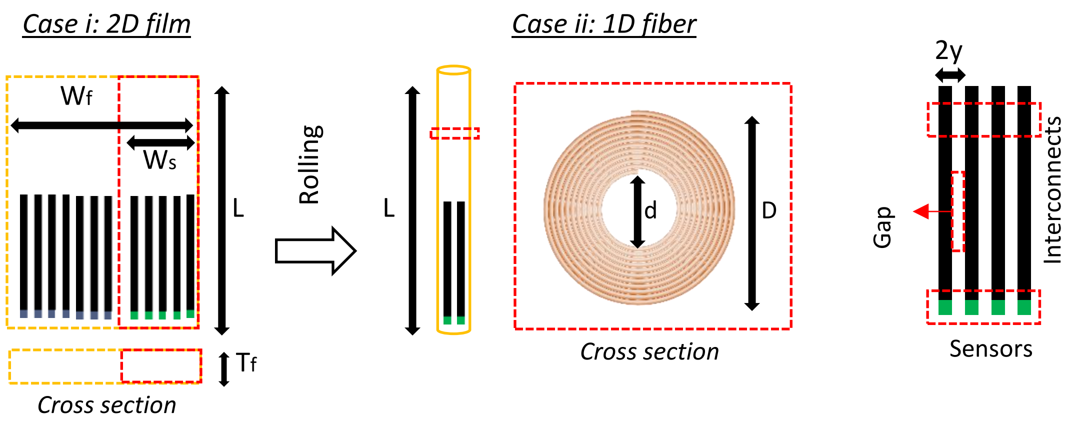


The following assumption are used for the evaluation:

1- Ideal rolling.

2- N is the total number of electronic components in the 2D film and fiber and is defined by the fabrication resolution (pitch size = 2y).

3- Internal diameter of the probe is equal to 5 times the thickness of the unrolled film (d = $5T_f$). We add this requirement to allow for easier calculations.

From the previous section:

$$(1) \qquad \frac{\pi L\,(D-d)^2}{4} = T_f W_f L$$

$$(2) \qquad W_f = \frac{\pi (D-d)^2}{4T_f}$$

Defining the density of electrical components on the planar 2D film ($\rho_f$):

$$(3) \qquad \rho_f = \frac{N}{W_f} = \frac{4T_f N}{\pi (D-d)^2}$$

Defining the density of electrical components in a rolled fiber (including both internal and surface sensors):

$$(5) \qquad \rho_r = \frac{N}{D}$$

Defining the density gain (G) and applying d = 5 $T_f$:

$$(6) \qquad G = \frac{\rho_r}{\rho_f} = \frac{\frac{N}{D}}{\frac{4\mathrm{T}_f N}{\pi(\mathrm{D-d})^2}} = \frac{\pi(\mathrm{D-d})^2}{4D\mathrm{T}_f} = \frac{\pi(\mathrm{D}-5\mathrm{T}_f)^2}{4D\mathrm{T}_f}$$

Examples: Considering the following range: D: 50 - 1000 µm and $T_f$: 1 - 50 µm.

(i) D = 1000 µm, d = 50 µm, and $T_f$ = 10 µm ➔ G = 71.0

(ii) D = 200 µm, d = 50 µm, and $T_f$ = 10 µm ➔ G = 8.8

(iii) D = 200 µm, d = 10 µm, and $T_f$ = 2 µm ➔ G = 70.8

(iv) D = 500 µm, d = 50 µm, and $T_f$ = 1 µm ➔ G = 317.9

For surface components, the simplest model will give the lowest possible gain: G = π**.**

**Supplementary Section S3.** A mathematical model describing the transformation of a 2D film into an Archimedean spiral.

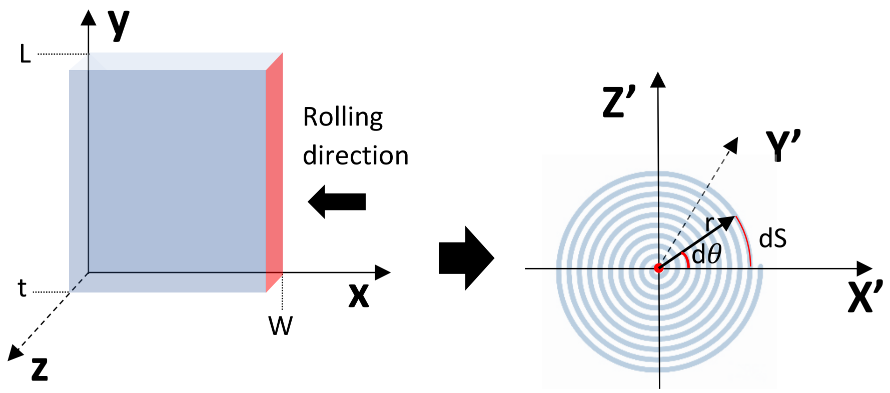


- $\mathrm{dS} = \sqrt{r^2 + \left(\frac{dr}{\mathrm{d}\theta}\right)^2}\, \mathrm{d}\theta$
- For this specific Archimedean spiral: r = a + b$\theta$, where a = 0, b = 2t/2$\pi$ (https://mathworld.wolfram.com/ArchimedesSpiral.html).
- After substitution:

$$\mathrm{dS} = \sqrt{(\frac{t}{\pi}\theta)^2 + \left(\frac{(\frac{t}{\pi})\mathrm{d}\theta}{\mathrm{d}\theta}\right)^2}\, \mathrm{d}\theta = \sqrt{(\frac{t}{\pi}\theta)^2 + \left(\frac{t}{\pi}\right)^2}\, \mathrm{d}\theta = \frac{t}{\pi}\sqrt{\theta^2 + 1}\, \mathrm{d}\theta$$

- After Integration:

$$\mathrm{S} = \frac{t}{2\pi}\left(\theta\sqrt{1+\theta^2} + sinh^{-1}(\theta)\right) = \frac{1}{2}\left(r\sqrt{1+\left(\frac{\pi r}{t}\right)^2} + sinh^{-1}(\frac{\pi r}{t})\right)$$

- The spiral transformation maps the (X,Y) coordinates of the 2D film into the following coordinates:
- X ➔ S = $\frac{1}{2}\left(\theta\sqrt{1+\theta^2} + sinh^{-1}(\theta)\right)$, Y ➔ Y'
- Using these equations, precise control over the angular, longitudinal, and radial positions can be achieved. **Y'** represents the fiber longitudinal axis. **S** can be used to determine the exact r and $\theta$.

### Supplementary Tables

**Table S1.** Comparison of the bioelectronic fibers developed in this work with other biomedical fibers from the literature.

| Technology | Fiber diameter | Estimated modulus (MPa) | Preparation method | Total number of channels | Functions | Limitations |
|---|---|---|---|---|---|---|
| Multifunctional brain probe [20] | 70 to 700 μm | 2380 | Thermal Drawing | 36 electrodes on a ~ 250 μm diameter fiber | Wave guidance, fluid delivery, and single neuron recording) | 1. Functions localized at the tip of the fiber<br>2. Temperature tolerance up to 325 °C<br>3. Microfabrication not shown |
| Fiber with integrated electronic components [21] | 150 μm | 70000 | Maskless photolithography on square-shaped microfibers | 30 | Ring oscillators, inverters, phototransistors, condensers, and temperature sensors | 1. Limited to square-shaped fibers<br>2. Microfabrication done on each fiber individually |
| Bioelectronic sutures for the monitoring of deep surgical wounds [22] | 400 μm | 3000 | Modification of commercial silk fibers | One channel per fiber | Detection of gastric fluids leakage | 1. low density<br>2. Low functionality<br>3. Did not show control over device position and orientation<br>4. Microfabrication not shown |
| Suturable fiber sensors for wireless monitoring of connective tissue strain [23] | 700 - 1300 μm | 3.5 | Modification of polyurethane-based commercial stretchable fibers | One channel per fiber | Strain sensing | 1. low density<br>2. Low functionality<br>3. Did not show control over device position and orientation<br>4. Microfabrication not shown |
| Multiply sensing fibers [24] | 150 μm | 118 | Helical bundle of functionalized carbon nanotubes fibres | 6 | Glucose, Ca2+, Na+, K+ and pH | 1. low density<br>2. Did not show control over device position and orientation<br>3. Microfabrication not shown |
| This work | 50 to 1200 μm | 0.2 – 81 | Microfabrication followed by 2D-to-1D dimensionality reduction | ~ 150 on a 250 μm fiber (demonstrated) | Serotonin, pressure, and pH sensing; electrical stimulation; single-neuron recording; fluid delivery; and light guiding | May need a dissolvable rigid coating or a guiding wire for fiber insertion if it is made of a highly soft substrate |

**Table S2.** Comparison of the bioelectronic fibers developed in this work with commercial products and probes from the literature used specifically for motility recording in the gastrointestinal tract.

| Technology | Probe diameter | Probe cost | Functions | Minimum sensor spacing | Total sensor numbers | Notable advantages | Notable disadvantages |
|---|---|---|---|---|---|---|---|
| Water perfused catheters (commercial) | 4.7 mm [25]<br>4.2 mm [26]<br>4.6 [28] | $500 - $750 [27] | Hydraulic sensing | 1-2 cm [26];<br>10 cm [27];<br>0.8 cm [28];<br>1.5 cm [29] | 36 [25, 29];<br>22 [26];<br>8 [27];<br>10 [28]; | 1. Low probe cost<br>2. High configurability (e.g., rearrangeable, spatial distributions, variable number of channels)<br>3. Great sensor durability | 1. Requiring highly experienced personnel<br>2. Poor temporal resolutions<br>3. Bulkiness and high cost of water perfusion system<br>4. Large intervals between the pressure sensors<br>5. Prone to artifacts (e.g., air bubbles) |
| Solid-state transducers (commercial) | 3.8 – 5.4 mm [26];<br>3.5 mm [27];<br>4 – 5.3 mm [28];<br>3.3 [31];<br>10.75 [32] | $17,000 – 28,000 [27] | Piezoelectric and capacitive sensing | 1 cm [25, 26, 30];<br>10 cm [27];<br>0.8 cm [28];<br>3 cm [29];<br>2.5 cm [31];<br>4 mm axially and 2mm radially [32] | 36 [25, 26, 29, 30];<br>8 [28];<br>40 [31]<br>256 [32] | 1. High spatiotemporal resolutions<br>2. Portability of sensing systems | 1. High overall cost<br>2. Sensor fragility |
| Optical fiber pressure sensors | 3 mm [33];<br>2.2 mm [34] | Unknown | Optical sensing | 1 cm [33, 34, 35, 36] | 32 [33];<br>72 [34];<br>72-90 [35];<br>72 [36]; | 1. High channel density due to thin fiber dimeters<br>2. Minimal invasiveness during placement | 1. Sensor Fragility<br>2. High material and fabrication barriers<br>3. Bulkiness and high cost of recording systems |
| Liquid metal-enabled pressure transducers (QUILT) [37] | 0.64 mm (one channel) | $ 0.26/cm | Resistive sensing | 0.5 cm | 8 (1 per fiber) | 1. Low cost<br>2. High spatiotemporal resolutions<br>3. Portability of recording systems | 1. Relatively large percentage uncertainty<br>2. Variations due to sensor orientations |
| This work | 50 to 1200 µm | Low cost | Piezoresistive and electrochemical sensing; Electrical stimulation | 300 µm | ~ 150 (on a 250 µm fiber) | 1. High density<br>2. Multifunctional pH, serotonin, and pressure sensing; electrical stimulation, fluid delivery, light guiding<br>3. Compatibility with conventional microfabrication | May need a dissolvable rigid coating or a guiding wire for fiber insertion if it is made of a highly soft substrate |

**Table S3.** Comparison between S-NeuroString and other existing technologies used for single neuron recording.

| Probe Type | Material | Young's Modulus (MPa) | Shank shape | Shank thickness (um) | Shank width (um) | Stiffness (GPa.um^4) | Total channel count | Channel per shank | Stretchable | Longest time recorded (weeks) |
|---|---|---|---|---|---|---|---|---|---|---|
| Thin-film [38] | Polyimide | 4000 | Rectangular | 3 | 12 | 108 | 16 | 1 | N | 6 |
| Microwire [39] | PtIr | 200000 | Cylinder | 15 | 15 | 497010 | 251 | 1 | N | Acute |
| Microwire [40] | Silicon | 168900 | Cylinder | 80 | 80 | 339593600 | 96 | 1 | N | 132 |
| Thin-film [41] | SU8-100 | 2200 | Rectangular | 1 | 30 | 5.5 | 1024 | 128 | N | 41 |
| Thin-film [42] | Polyimide | 4000 | Rectangular | 14 | 80 | 73173 | 1024 | 16 | N | 23 |
| Thin-film [43] | SU8 2000.5 | 2000 | Rectangular | 0.9 | 2 | 0.243 | 16 | 1 | N | 13 |
| Thin-film [44] | Parylene-C | 2800 | Rectangular | 4 | 75 | 1120 | 14 | 14 | N | 1 |
| Thin-film [45] | Polyimide | 4000 | Rectangular | 4 | 50 | 747 | 3072 | 32 | N | Acute |
| Microwire [46] | PtIr | 200000 | Cylinder | 18 | 18 | 1030599 | 30000 | 1 | N | Acute |
| Fiber [20] | Polycarbonate | 2380 | Cylinder | 416 | 416 | 3498808543 | 36 | 36 | N | 8 |
| Fiber [20] | Polycarbonate | 2380 | Cylinder | 85 | 85 | 6098493 | 7 | 7 | N | 8 |
| Fiber [47] | Borosilicate | 64000 | Cylinder | 25 | 25 | 1227185 | 128 | 1 | N | Acute |
| Fiber [48] | Carbon fiber | 241000 | Cylinder | 6.8 | 6.8 | 25294 | 16 | 1 | N | 12 |
| Fiber [49] | Carbon | 255000 | Cylinder | 6.8 | 6.8 | 26764 | 16 | 1 | N | 12 |
| Microwire [50] | Silicon | 168900 | Cylinder | 10 | 10 | 82909 | 1024 | 1 | N | 28 |
| Fiber [51] | Polyurethane/PDMS | 0.17 | Cylinder | 172 | 87 | 6199 | 1 | 1 | Y | 16 |
| Fiber [52] | Hydrogel coated - poly(etherimide) | 3300 | Cylinder | 334 | 334 | 2015903347 | 21 | 21 | N | 24 |
| Shank [53] | Silicon | 168900 | Rectangular | 24 | 70 | 2750708 | 10,240 | 1280 | N | 44 |
| Shank [54] | Silicon | 168900 | Rectangular | 50 | 100 | 51817871 | 1344 | 1344 | N | Acute |
| This work (fiber) | SBS | 81.2 | Cylinder | 150 | 150 | 2017860 | 32 | 32 | Y | 2 |

### Supplementary Movies

**Supplementary Movie S1|** The transformation process of a 3 cm wide and 12 cm long film into a 300 μm fiber. Click on the following link to download the movie:

**Supplementary Movie S1**
**Click here to download**
**Supplementary Movie S1**

**Supplementary Movie S2|** Ex-vivo detection of progressive contractions using S-NeuroString. Click on the following link to download the movie:

**Supplementary Movie S2**
**Click here to download**
**Supplementary Movie S2**

**Supplementary Movie S3|** The movie shows the bioelectronic fiber inserted in the small intestine of pig and used to record the observed natural motility patterns. Click on the following link to download the movie:

**Supplementary Movie S3**
**Click here to download**
**Supplementary Movie S3**

**Supplementary Movie S4|** The movie shows the programmable stimulation system which allows the stimulation of specific tissue part. Click on the following link to download the movie:

**Supplementary Movie S4**
**Click here to download**
**Supplementary Movie S4**
